# Modelling oxaliplatin resistance in colorectal cancer reveals a *SERPINE1*-based gene signature (RESIST-M) and therapeutic strategies for pro-metastatic CMS4 subtype

**DOI:** 10.1101/2024.12.17.628817

**Authors:** Stephen Qi Rong Wong, Mohua Das, Niranjan Shirgaonkar, Kenzom Tenzin, Huiwen Chua, Lin Xuan Chee, Sae Yeoh Ahpa, Astley Aruna Murugiah, Wei Yong Chua, Madelaine Skolastika Theardy, Matan Thangavelu Thangavelu, Jane Vin Chan, Choon Kong Yap, Iain Bee Huat Tan, Petros Tsantoulis, Sabine Tejpar, Jia Min Loo, Ramanuj Dasgupta

## Abstract

Drug resistance and distant metastases are major contributors to mortality in colorectal cancer (CRC). Here we investigate mechanisms underlying acquired resistance to oxaliplatin, a first-line, standard-of-care CRC treatment. We generated oxaliplatin-resistant CRC tumor cells with clinically relevant dosing regimen, which displayed enhanced metastatic potential. Transcriptomic and phenotypic analyses revealed a critical function for cholesterol biogenesis in modulating TGF-β signaling activity, which in turn regulates *SERPINE1* expression, a gene we identified as a key player in promoting drug resistance and metastasis. Additionally, we uncovered a *SERPINE1*-associated nine-gene expression signature, RESIST-M, that can predict overall and relapse-free survival (RFS) in clinical cohorts and is able to stratify patients into CMS4/iCMS3-fibrotic CRC-subtypes, underscoring its clinical utility. Using mouse tumor models, we provide further evidence that targeting *SERPINE1* and cholesterol biogenesis can be viable approaches to re-sensitize the resistant pro-metastatic CRC cells to oxaliplatin. This study not only elucidates the molecular underpinnings of drug resistance and metastasis in primary CRC, but also offers prognostic and therapeutic strategies to guide clinical management of the disease.

**Significance:** This study reveals critical resources and insights on oxaliplatin resistance and metastasis in CRC via a novel TGF-β cholesterol axis. We generated improved oxaliplatin-resistant models that enabled identification of a prognostic *SERPINE1*-based gene signature to predict oxaliplatin resistance-induced metastasis in CRC. This gene signature derived from our models showed that the models can mimic CMS-4/iCMS-fibrotic-like metastatic CRC patients. We validated therapeutic candidates targeting CMS-4/iCMS-fibrotic-like metastatic CRC cells which can reverse drug resistance and metastasis.

## Introduction

Colorectal cancer (CRC) is a leading cause of cancer mortality globally. While over 25% of CRC patients present themselves with distant metastatic disease at diagnosis, ∼40-50% of patients will eventually develop metastases over the course of neoadjuvant or post-surgical adjuvant therapy^1,2^. This suggests the presence of prior disseminated cancer cells in distal sites, primarily in the liver or lungs, that are either resistant to chemotherapy or acquire resistance during the course of treatment. The 5-year survival rate for patients with unresectable metastatic CRC remains a dismal 15%^1,2^, highlighting the need to identify molecular mechanisms driving therapeutic resistance and metastatic relapse.

Despite advancements in targeted and immunotherapy, combinatorial adjuvant chemotherapy, such as FOLFOX and FOLFIRI, remains the cornerstone of CRC treatment. The MOSAIC^3^ and NSABP C-07^4^ clinical trials established oxaliplatin as a standard adjuvant treatment for resected stage II/III colon cancer. These trials demonstrated that adding oxaliplatin to the regimen significantly improves disease-free survival (DFS) when combined with 5-fluorouracil (5-FU) and leucovorin (LV)^3,4^. However, follow-up reports from the MOSAIC trial have indicated that the improvement in absolute overall survival (OS) for resected stage III CRC patients with the addition of oxaliplatin was only 4.6% over 10 years. Notably, patients with the stem-like CMS-4 subtype of CRC, which is linked to poor prognoses, derived the least benefit from oxaliplatin in the NSABP C-07 and MOSAIC trials^5^. Therefore, it remains critical to identify the subset of patients with stage II/III disease who are likely to relapse and more importantly, likely to benefit from oxaliplatin.

Mechanisms of tumor cell resistance to oxaliplatin have been extensively studied^6^. Recent research has highlighted the role of cell cycle-dependent kinases (CDK1)^7^, pyruvate kinase M2 (PKM2)^8^, and drug transporters^9^ as key mediators of acquired resistance to oxaliplatin. Additionally, the activation of CD95 has been implicated in enhancing the metastatic potential of oxaliplatin-resistant cells by promoting epithelial-to-mesenchymal transition (EMT) characteristics^10^. Notably, inhibiting glutathione synthesis has been shown to sensitize peritoneal metastases in colorectal cancer patient-derived organoids (PDOs) to oxaliplatin treatment^11^, emphasizing the importance of targeting acquired vulnerabilities in oxaliplatin-resistant metastatic cancers. Despite these advancements, a systematic and clinically relevant understanding of how oxaliplatin-resistant colorectal cancer cells develop metastatic properties and impact patient prognosis remains relatively underexplored in the current literature.

In this study, we modelled CRC progression to metastatic disease following treatment by exposing two CRC cells lines to clinically relevant doses of oxaliplatin. Molecular and phenotypic characterization revealed the activation of a cholesterol-TGF-β-*SERPINE1* signaling axis during treatment that appears to promote oxaliplatin resistance and enhanced metastatic potential in one of the cell lines, HCT116. Additionally, a nine-gene expression signature, RESIST-M, was identified, stratifying patients with the CMS4 subtype as well as a high-risk iCMS3-MSS-fibrotic subtype associated with the poorest overall survival. We observed that RESIST-M signature could specifically distinguish CMS4 CRC and hence predict patient survival better than previously reported gene signatures of oxaliplatin resistance, Finally, we discovered specific vulnerabilities in oxaliplatin-resistant CRCs, presenting new therapeutic opportunities. Our findings suggest promising pathways for interventions aimed at curbing metastatic progression and reducing mortality among oxaliplatin-resistant CRC patients, particularly those with the CMS4/iCMS3-MSS-fibrotic subtype.

## Results

### Clinically relevant dosing of CRC cell lines induced a drug-resistant and metastatic phenotype

Previous studies on chemoresistance and metastasis often used models with prolonged or supra-physiological exposure to drugs, which is not reflective of clinical conditions^12,13^. It is therefore crucial to investigate cellular changes under clinically relevant therapeutic exposure. To evaluate the effects of oxaliplatin, we subjected two CRC cell lines, HCT116 and SW480, to a regime consisting of 10 treatment cycles at three different concentrations: 0.5 µM (Low-Dose; [HCT116-LD or SW480-LD]), 5 µM (Mid-Dose; [HCT116-MD or SW480-MD]) and 80 µM (High-Dose; [HCT116-HD or SW480-HD]) (**Fig. 1A**), corresponding to 0.1x, 1x and 16x Cmax observed in patients^14^.

**Figure 1.**
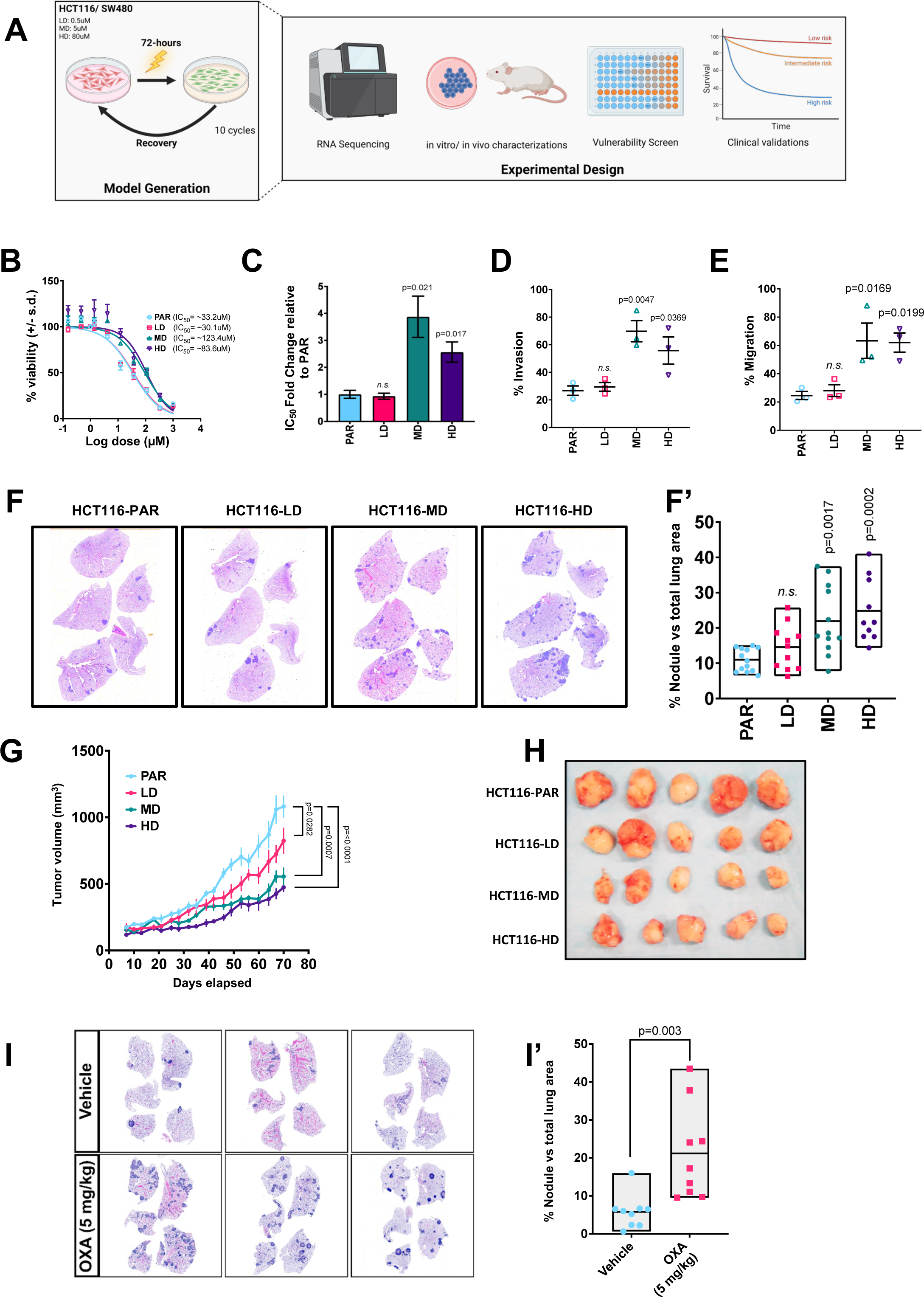
Cyclical dosing of oxaliplatin in CRC cell lines induced a drug-resistant and metastatic phenotype. A, Scheme depicting generation of oxaliplatin resistant models. Parental cells (PAR), either SW480 or HCT116, were treated with 0.5µM (LD), 5µM (MD) or 80µM (HD) of oxaliplatin for 72hrs and then recovered in drug-free medium until confluent. This dosing regime was repeated for 10 cycles. Developed oxaliplatin resistant models were subjected to downstream transcriptomic and phenotypic characterizations compared to parental cells. B-E, *In vitro* characterization of oxaliplatin resistant HCT116 models. Representative dose-response curves of oxaliplatin resistant models compared to parental cells (**B**). HCT116 parental (PAR), low dose (LD), mid dose (MD) and high dose (HD) were incubated with oxaliplatin for 72 hrs followed by measurement of IC_50_. Fold change in IC_50_ of oxaliplatin resistant models compared to parental cells (**C**). Fold changes are shown for HCT116 (LD, MD and HD) relative to PAR from three sets of independent dose-response IC_50_ studies. Transwell invasion assay in oxaliplatin resistant HCT116 models (LD, MD and HD) and parental cells (PAR) (**D**). Percentage invasion was determined by measuring the number of cells that had invaded to the bottom chamber divided by total number of cells seeded. Transwell migration assay in oxaliplatin resistant HCT116 models (LD, MD and HD) and parental cells (PAR) (**E**). Percentage migration was determined by measuring the number of cells that had invaded to the bottom chamber divided by total number of cells seeded. F-H, *In vivo* characterization of oxaliplatin resistant HCT116 models. Representative H&E staining and corresponding quantifications of lung sections (**F, F’**) in mice bearing tumors of oxaliplatin resistant and parental HCT116 cells (refer to section ‘Tumor growth kinetics and spontaneous metastasis’ under Materials and Methods). Quantification of nodule positive area vs total lung area in mice harboring HCT116 PAR, LD, MD and HD xenografts (n= at least 8). Tumor growth kinetics *in vivo* of HCT116 PAR, LD, MD and HD xenografts (n= 5) (**G**). Corresponding images of tumor of HCT116 PAR, LD, MD and HD xenograft at day 70 (**H**). **I-I’, Spontaneous metastasis model using oxaliplatin treated HCT116.** Three representative H&E staining and corresponding quantifications (**I and I’**) of lung sections from mice harboring HCT116 PAR xenografts treated with vehicle or oxaliplatin for spontaneous metastasis (refer to section ‘*In vivo* effects of oxaliplatin treatment on spontaneous metastasis’ under Materials and Methods). Quantification of nodule positive area vs total lung area in mice harboring HCT116 PAR, LD, MD and HD xenografts (n= 9). *In vivo* effects of oxaliplatin treatment on spontaneous metastasis. Statistical significance was determined using Ordinary one-way ANOVA followed by Dunnett’s multiple comparisons test for Figs **C-F’**, using Ordinary two-way ANOVA followed by Dunnett’s multiple comparisons test for Fig **G**, and using two-tailed unpaired t-test for Fig **I’**. A p-value of <0.05 was considered significant for all analyses, unless stated otherwise.

After 10 cycles of treatment, we observed that cells treated at MD and HD (i.e., 5 µM and 80 µM), but not LD (i.e., 0.5 µM), displayed increased resistance to oxaliplatin (**Fig. 1B, C; Supplementary Fig. S1A, B**). We also observed a “spindle-like” morphology in HCT116-HD cells, not observed in other treatment arms (**Supplementary Fig. S1C**), suggesting high dose treatment may induce EMT-like morphology. Intriguingly, oxaliplatin-treated HCT116-MD and HD, but not SW480-MD/HD, also gained migratory and invasive capabilities associated with metastatic phenotypes *in vitro* (**Fig. 1D, E; Supplementary Fig. S1D, E)**. Additionally, HCT116-MD and HD cells displayed enhanced spontaneous lung metastases in immune deficient mouse xenograft models, *in vivo* (**Fig. 1F; Supplementary Fig. S1F, G**), in agreement with the increased invasive characteristics observed *in vitro*. Notably, proliferation assays *in vitro* and *in vivo* demonstrated that the metastatic phenotype observed was not concordant with tumor cell-proliferation or their growth rates, with the HCT116-MD and HD cells displaying slower tumor-xenograft growth rates compared to the LD and untreated (parental) controls (**Fig. 1G, H; Supplementary Fig. S1H-K**).

To determine whether our findings are relevant *in vivo*, naïve HCT116 or SW480 tumor bearing mice were subjected to five cycles of oxaliplatin treatment (5 mg/kg). Corroborating our prior findings, oxaliplatin-treated mice harboring HCT116 xenografts demonstrated increased incidences of pulmonary metastasis that were not observed in mice with SW480 tumors (**Fig 1I; Supplementary 1L, M**). Owing to the strikingly increased potential of drug resistant HCT116 to metastasize, we decided to focus on this model for further investigations to probe the underlying mechanisms for drug-induced metastatic progression.

### *SERPINE1* is upregulated and promotes chemoresistance and metastasis in oxaliplatin-treated HCT116

In order to uncover potential molecular mechanisms driving the observed drug-resistant and metastatic phenotypes, we performed bulk RNA sequencing on oxaliplatin-treated and untreated cells. Gene Set Enrichment Analyses (GSEA) revealed up-regulation of well-established pathways consistent with a metastatic phenotype in HCT116-LD and MD cells such as EMT and angiogenesis (**Fig. 2A**), which was not observed in the SW480 cell lines (**Supplementary Fig. S2A, B**). Interestingly, we observed remarkably similar hallmarks being enriched between HCT116-LD and MD (**Supplementary Fig. S2A, B**), with unique pathways such as coagulation pathway and cholesterol homoeostasis, which we speculated could also contribute to drug-resistance and metastatic progression. Notably, gene sets enriched in HCT116-HD cells exposed to the supra-physiological concentration of 80 µM were markedly different from those observed with the low and mid doses (**Supplementary Fig. S2A**). This suggests that although cells exhibit similar phenotypes (i.e. drug resistance and increased metastasis), high doses of oxaliplatin might be selecting for cell-state dependent biological programs that allow survival of pro-metastatic drug-tolerant persistent cells.

**Figure 2.**
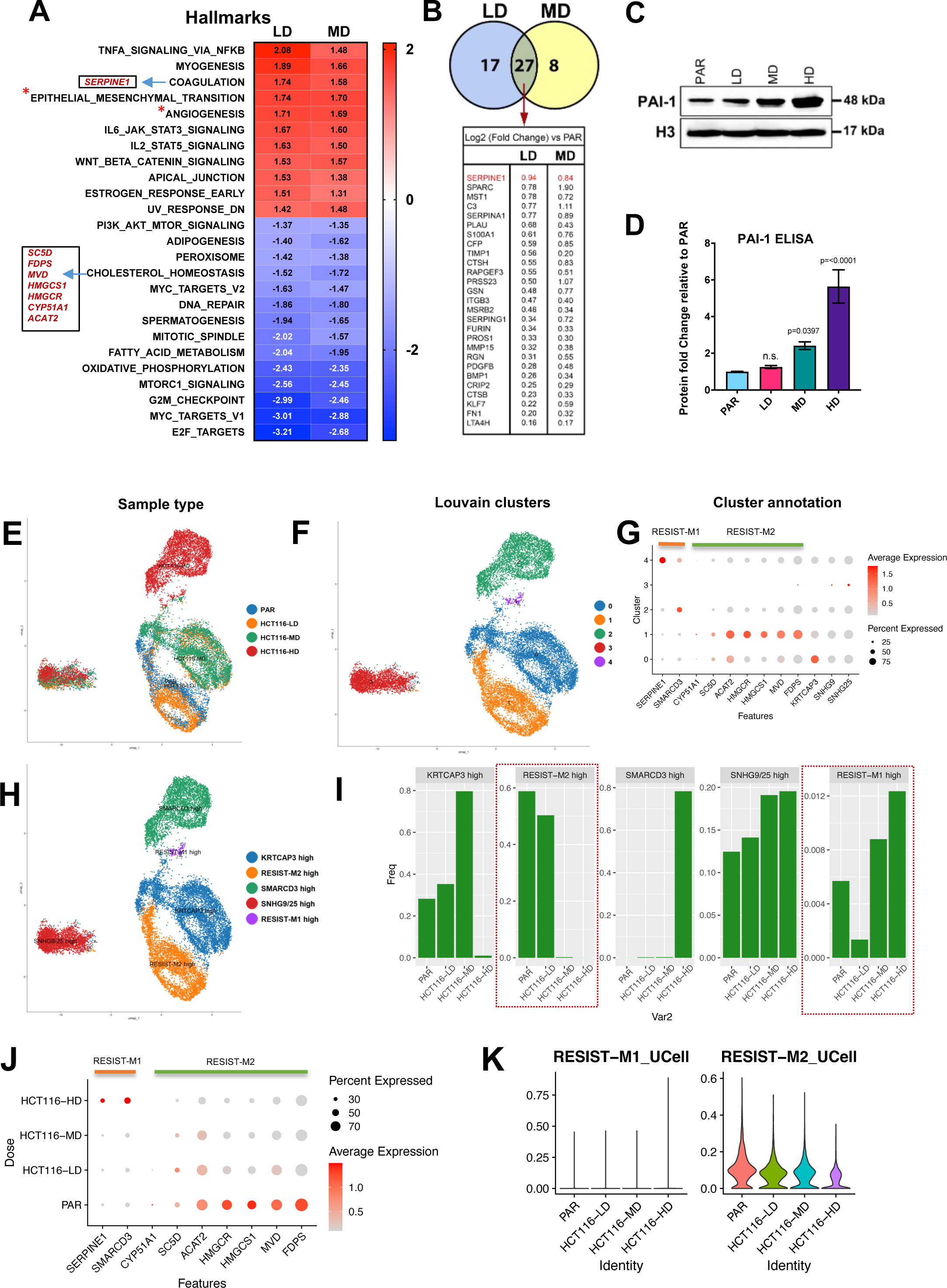
Overexpressed *SERPINE1* promotes chemoresistance and metastasis in oxaliplatin-treated HCT116. A-B, Bulk transcriptomic analyses of oxaliplatin resistant HCT116 models. Biological pathways enriched using differentially expressed genes (DEGs) in HCT116 LD/MD vs PAR (**A**). Red and blue indicates positively and negatively regulated genes respectively, with the values of normalized enrichment score (NES) as indicated. All hallmarks showed here are significantly enriched. Pathways with * symbol denotes pathways that are known to be hallmarks of drug-resistance and metastasis. Pathways and genes explored in this manuscript are highlighted with the gene names in black box. Differentially expressed genes (DEGs) in HCT116 LD/MD vs PAR (**B**). Fold changes in expression of each gene are indicated. C-D, Validation of enhanced *SERPINE1* (PAI-1) in oxaliplatin resistant HCT116 models. Immunoblotting of PAI-1 in HCT116 PAR, LD, MD and HD (**C**). Fold change of PAI-1 protein in cell culture supernatant of HCT116 LD, MD and HD relative to PAR determined using ELISA (**D**). **E-K, Single-cell transcriptomic analyses uncovers gene signature linked to oxaliplatin resistance (RESIST-M).** UMAP projection and sample type annotation of scRNA-seq data from the four scRNA-seq libraries: HCT116-PAR/LD/MD/HD (**E**). UMAP projection and Louvain-based clustering of the four samples identifies 5 distinct cell states (cl 0-5) defining the parental and resistant conditions (**F**). Dot plot showing differentially expressed genes (DEGs) of the 5 identified clusters (cl 0-5). RESIST-M gene signature, wherein RESIST-M1 genes denote *SERPINE1*, *SMARCD3*, and RESIST-M2 genes denote *CYP51A1*, *SC5D*, *ACAT2*, *HMGCR*, *HMGCS1*, *MVD*, *FDPS* can distinguish the cell states defined by clusters 1,2,4 (**G**). UMAP projection and annotation of the 5 identified cell states based on DEGs (**H**). Proportions of the different annotated clusters across the sample type shows loss and gain in clusters with high RESIST-M2 and high RESIST-M1 signature, respectively (**I**). Dot plot showing differential expression of RESIST-M genes as they increase and decrease respectively, depending on the dose of oxaliplatin and type of metastatic phenotype acquired (**J**). Violin plot showing Ucell score for RESIST-M1 and M2 genes, increasing and decreasing respectively, as the selection pressure of oxaliplatin is increased (**K**). Statistical significance was determined using Ordinary one-way ANOVA followed by Dunnett’s multiple comparisons test for Fig. **D.** A p-value of <0.05 was considered significant for all analyses, unless stated otherwise.

Amongst the enriched hallmarks in HCT116-LD and MD cells, we found a striking enrichment of the coagulation pathway. While prior research has implicated the coagulation pathways in tumor progression and metastatic spread^15,16^, specific molecular mechanisms remain relatively understudied. Overlapping genes that are differentially expressed from the coagulation hallmark between HCT116-LD and MD revealed *SERPINE1* as the top upregulated, and potentially druggable target gene (**Fig. 2B**). *SERPINE1* encodes the plasminogen activator inhibitor type 1 protein (PAI-1) and is known to play a critical function in maintaining homeostasis between fibrinolysis and thrombosis^17^. The protein has been implicated in drug resistance and metastasis in various cancer types^18,19^. Intriguingly, ELISA and immunoblotting of PAI-1 revealed upregulation only in HCT116-MD and HCT116-HD, but not in HCT116-LD (**Fig. 2C, D**), alluding to its potential involvement in acquired resistance and/or metastatic phenotypes.

To investigate therapy-induced acquired resistance and metastatic phenotypes at a more granular level, we performed single-cell RNA-sequencing (scRNA-seq) using the parental and resistant cell line models. Louvain-based clustering revealed five different cell states in the single-cell data (**Fig. 2E-I**). Amongst them, cluster 3 represented a unique cell state highlighted by the extensive loss of lineage-defining markers, with a few cells expressing *SNHG* (*SNHG9/25* high, *cl 3*). Notably, we found cl 3 to be shared between the parental and resistant cell lines (**Fig. 2E-I, Supplementary Fig. S2C**), suggesting the presence of pre-existing resistant cell-states that get selected upon oxaliplatin treatment. Additionally, using the genes identified from our bulk-RNA seq analyses, we uncovered two gene signatures, RESIST-M1 (where RESIST-M denotes **<u>RESI</u>**stance-**<u>I</u>**nduced **<u>S</u>**igna**<u>T</u>**ure for CRC **<u>M</u>**etastasis) defined by high levels of *SERPINE1* and *SMARCD3* expression; and RESIST-M2, defined by the expression of *CYP51A1*, *SC5D*, *ACAT2*, *HMGCR*, *HMGCS1*, *MVD*, *FDPS*. As shown in **Fig. 2G**, the RESIST-signatures could distinguish two distinct cell states represented by cl 4 (RESIST-M1) and cl 1 (RESIST-M2). We observed that the RESIST-M1 and RESIST-M2 signatures were gradually gained and lost, respectively, as the cells became more resistant and gained metastatic potential (**Fig. 2H-I, 2J-K**). For example, oxaliplatin resistance correlated with the gain in RESIST-M1 and *SMARCD3* expression, which also exhibited increased *SERPINE1* expression (**Supplementary Fig. S2D, E**). In contrast these clusters displayed a concomitant loss of RESIST-M2 signatures (**Fig. 2H-I**). Intriguingly, a distinct, divergent evolutionary trajectory was observed for HCT116-HD cells that showed high *SMARCD3* expression, a concomitant loss of *KRTCAP3* expression, and the loss of heterogeneity, as defined by the predominance of a single cluster, cl 2 (**Fig. 2I**, middle panel). This loss of heterogeneity observed in cells selected with high dose oxaliplatin was also accompanied by a gradual increase in differentiation markers, as revealed by cytotrace analysis (**Supplementary Fig. S2F-H**). Pseudotime analyses, excluding *SNHG9/25* high cluster, indeed demonstrated that HCT116-HD cell lines emerge as a distinct, adaptive drug-tolerant cell population (**Supplementary Fig. S2I-J**). In contrast, cell populations treated with clinically relevant concentrations (LD and MD) revealed more of a continuum of cell-state trajectories, which partly retained transcriptomic heterogeneity found in the parental, untreated cells (**Fig. 2E-H, Supplementary Fig. S2I-J**). Taken together, these data suggest that both selection of distinct cell-states defined by their genomic and/or transcriptomic heterogeneity, as well as adaptive alterations in gene expression upon distinct dosing schedules, can drive distinct molecular trajectories and establishment of phenotypic states.

### Genetic and pharmacological inhibition of *SERPINE1* reverses chemoresistance and metastasis in oxaliplatin-treated HCT116

To investigate a functional role of PAI-1, we conducted loss-of-function studies using two distinct shRNAs targeting the *SERPINE1* gene (**Fig. 3A-C**). Notably, shRNA mediated knockdown of PAI-1 re-sensitized HCT116-MD and HCT116-HD to oxaliplatin treatment (**Fig. 3D-E**). Importantly, pharmacological inhibition of PAI-1 using tiplaxtinin^20^ resulted in significantly lower incidences of spontaneous pulmonary metastases in HCT116 tumor bearing mice treated with oxaliplatin (**Fig. 3F,F’**). Our findings thus far demonstrated a previously unrecognized role of oxaliplatin-induced *SERPINE1* in promoting chemoresistance and metastasis in colorectal cancer.

**Figure 3.**
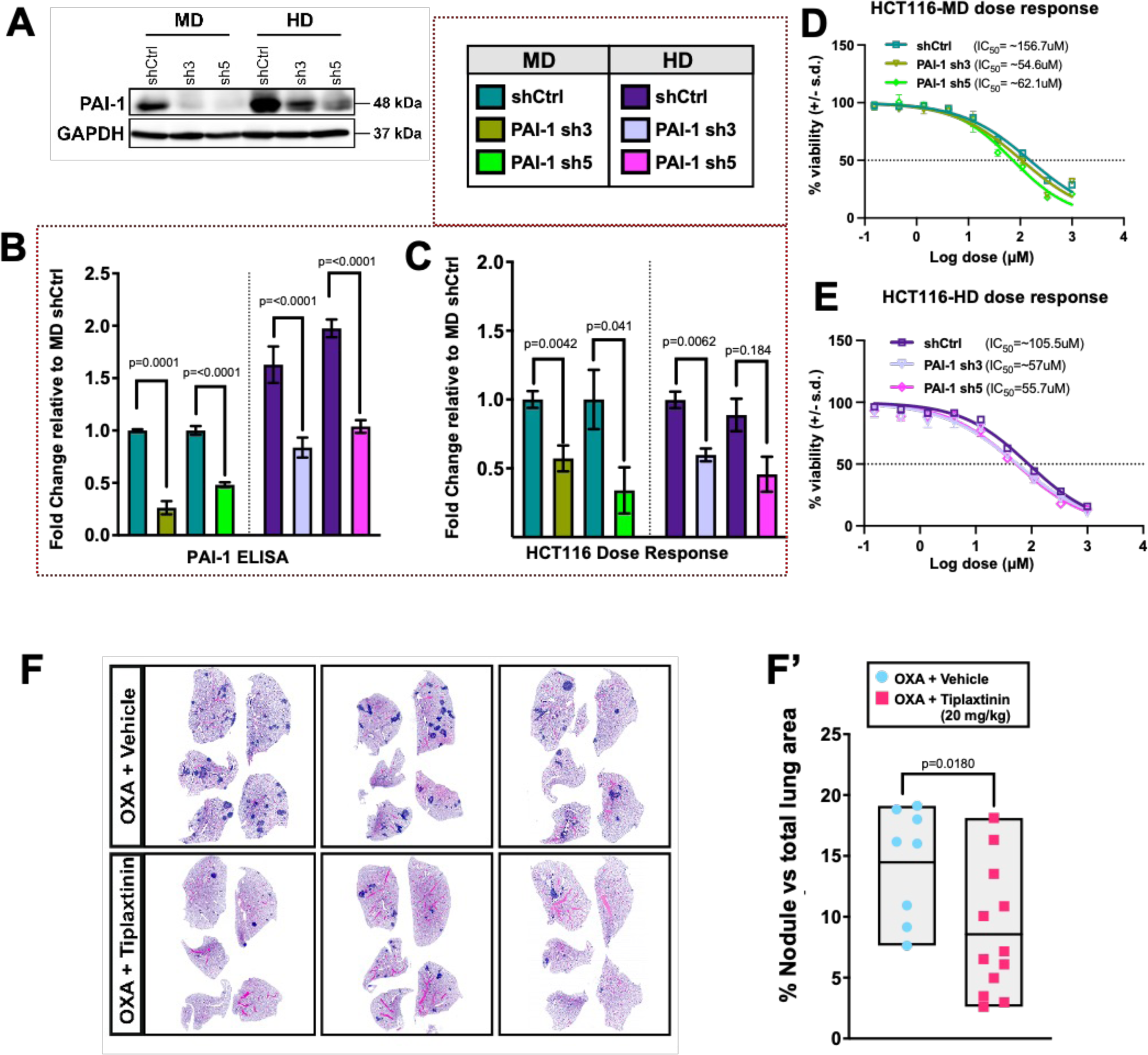
Genetic and pharmacological inhibition of *SERPINE1* can reverse chemoresistance and metastasis in oxaliplatin-treated HCT116. A-E, Genetic knockdown of *SERPINE1* in oxaliplatin resistant HCT116 models. Immunoblotting of PAI-1 protein in HCT116 MD and HD knocked down for the protein using two unique shRNAs (**A**). Fold change of PAI-1 protein in cell culture supernatant of HCT116 MD and HD knocked down for the protein relative to MD shRNA control (shCtrl) (**B**). Fold change of dose response to 48 hrs oxaliplatin treatment in HCT116 MD and HD knocked down for the protein relative to MD shCtrl (**C**). Representative dose response curves of HCT116-MD shCtrl, PAI-1 sh3 and sh5 (**D**) and HCT116-HD shCtrl, PAI-1 sh3 and sh5 (**E**) to 48 hrs of oxaliplatin treatment. **F-F’, Inhibition of metastasis in oxaliplatin treated HCT116 model using tiplaxtinin.** Three representative H&E staining and corresponding quantifications (**F and F’**) of lung sections from mice harboring HCT116 PAR xenografts treated with oxaliplatin, alone or in combination with tiplaxtinin, until spontaneous metastasis (refer to section ‘Impact of concomitant administration of tiplaxtinin and oxaliplatin on metastasis’ under Materials and Methods). For quantification number of nodule positive area was normalized to total lung area (n= at least 8). Statistical significance was determined using Ordinary one-way ANOVA followed by Sidak’s multiple comparisons test for Fig. B, C, and using two-tailed unpaired t-test for Fig. **F’.** A p-value of <0.05 was considered significant for all analyses, unless stated otherwise.

### *SERPINE1* upregulation is associated with dysregulated cholesterol-TGF-β signaling

Given *SERPINE1’s* pivotal role in facilitating metastasis and resistance in our investigations, we interrogated potential molecular mechanisms responsible for its upregulation following oxaliplatin treatment. Incidentally, *SERPINE1* is known to be transcriptionally regulated by TGF-β signaling^21^. Immunoblot analysis of key proteins involved in the TGF-β signaling pathway revealed elevated levels of this cytokine (TGF-β) and increased canonical SMAD2/3 activity in both HCT116-MD and HD cell lines (**Fig. 4A, B**). These results suggest that oxaliplatin treatment augments canonical TGF-β signaling, thereby promoting PAI-1 expression in HCT116 cells.

**Figure 4.**
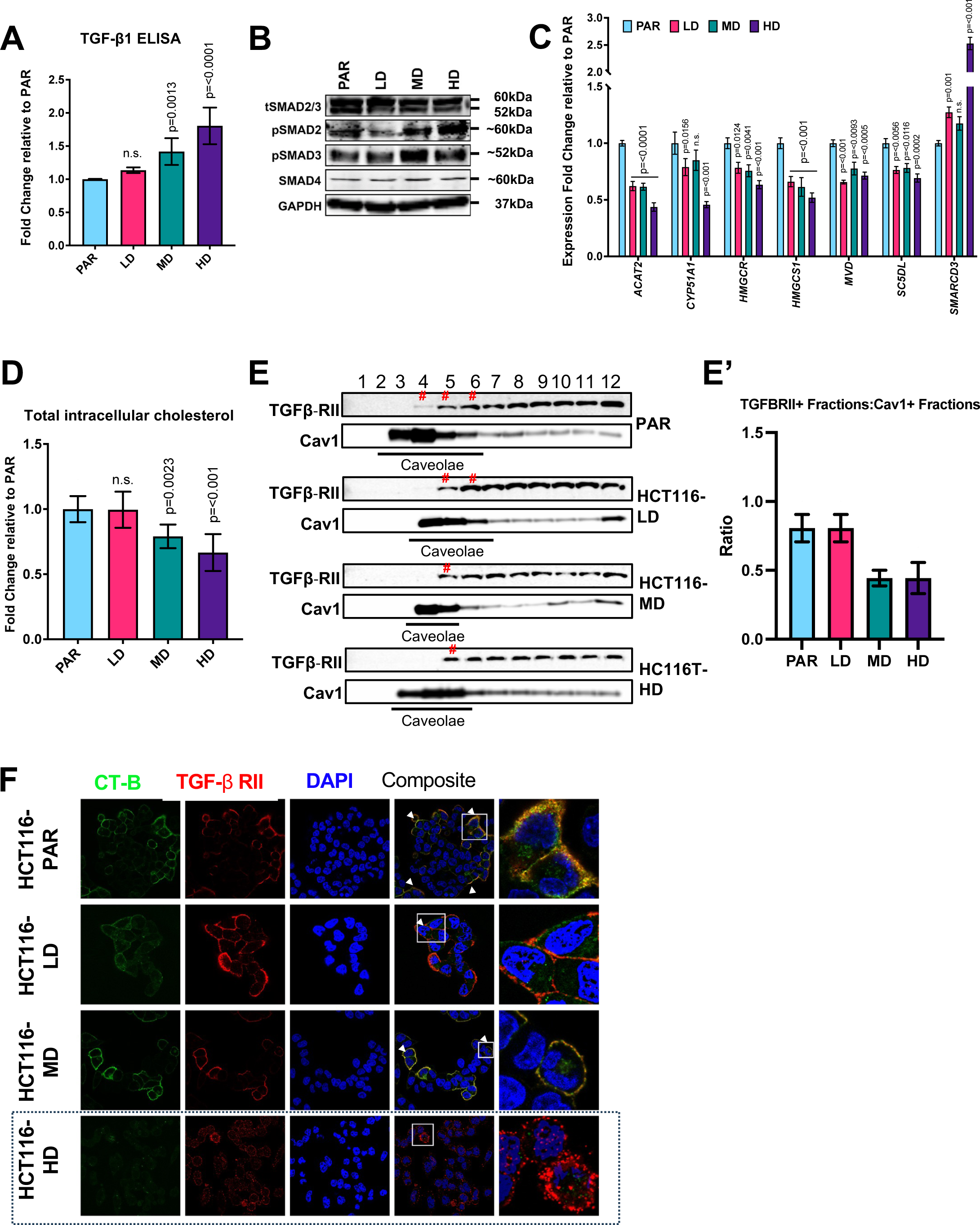
*SERPINE1* upregulation is associated with dysregulated cholesterol-TGF-β signaling. A-B, TGFb signaling is dysregulated in oxaliplatin resistant HCT116 models. Fold change of TGF-β1 protein in cell culture supernatant of HCT116 LD, MD and HD relative to PAR determined using ELISA (**A**). Immunoblotting of total SMAD2/3 (tSMAD2/3), phospho-SMAD2 (pSMAD2), phospho-SMAD3 (pSMAD3) and SMAD4 in oxaliplatin resistant HCT116 models (**B**). **C-F, Altered cholesterol biogenesis and reduction of caveolae-coated TGFbRII characterizes oxaliplatin resistant HCT116 models.** Validation of cholesterol-related genes identified in Fig. 2 with their fold changes in the resistant models relative to parental cells determined using quantitative real-time PCR (**C**). Fold change of total intracellular cholesterol in resistant models relative to parental cells determined using ELISA (**D**). Representative immunoblot of TGFBRII and caveolin after fractionation. # indicates TGFBRII found in lipid raft (caveolin-high/positive) fractions (**E**). Quantification of three independent immunoblots denoting the ratio of TGFBRII positive fractions to Cav1 positive fractions (**E’**). Immunofluorescence (IF) staining for colocalization of cholera toxin subunit B (CT-B; green indicating lipid rafts) and TGF-β receptor II (TGF-β RII; red) (**F**). White arrows indicate TGFβRII internalized in lipid rafts. White box denotes selected regions that are magnified. Black dotted box highlights the high oxaliplatin dose (HCT116 HD) that show most drug-tolerant persistent cells with reduced lipid rafts colocalized with TGFβRII. Statistical significance was determined using Ordinary one-way ANOVA followed by Dunnett’s multiple comparisons test for Fig. A, D, and using Ordinary 2way ANOVA followed by Dunnett’s multiple comparisons test for Fig. **C.** A p-value of <0.05 was considered significant for all analyses, unless stated otherwise.

TGF-β signaling is tightly regulated through various mechanisms, including receptor endocytosis localized in distinct compartments of the plasma membrane, such as cholesterol-rich lipid rafts^22^. Most notably, GSEA analysis of RNA-seq data from oxaliplatin-treated HCT116 cells indicated a significant decrease in genes involved in the cholesterol biosynthesis pathway (**Fig. 2A, 4C**). Specifically, we identified eight genes significantly enriched in processes directly and/or indirectly related to cholesterol biogenesis (**Supplementary Fig. S3A**) and further validated seven of them using qPCR (**Fig. 4C**). Additionally, measurement of intracellular cholesterol levels revealed a significant reduction in sterol content in HCT116-MD and HD cells (**Fig. 4D**).

Cholesterol is an essential component of lipid-rafts in the cell’s plasma membrane, and these specialized compartments have been known to play a key role in modulating various signaling cascades by controlling the localization of receptors between raft and non-raft regions^23^. Given the observed decrease in cholesterol biosynthesis in HCT116-MD and HD cells, we hypothesized that this could impact TGF-β signaling by altering the distribution of TGF-β receptors across the different membrane compartments (i.e. rafts and non-rafts regions). To investigate this, we employed sucrose density fractionation and immunofluorescence staining to examine the localization of TGF-β receptor 2 (TGFBRII) along the cell membrane. Remarkably, fractionation experiments demonstrated reduced amounts of TGFBRII within lipid raft domains of HCT116-MD and HD (**Fig. 4E, E’**). Consistently, immunofluorescence staining corroborated these findings, showing decreased colocalization of the receptors in lipid raft regions of the HCT116-MD and HD cells (**Fig. 4F**).

Our findings thus far indicate a potential association between lowered cholesterol levels and diminished localization of TGFBRII within lipid raft domains. It is known that TGF-β receptors localized in lipid rafts are more susceptible to ubiquitination and subsequent degradation^24^. In our investigation, the reduced presence of these receptors in lipid rafts suggests a plausible mechanism that enhanced signaling might occur because fewer receptors are targeted for degradation, potentially leading to increased SMAD2/3 activity that can further drive increased expression of PAI-1.

### RESIST-M signature significantly correlates with CMS4/iCMS3-fibrotic CRC subtype and predicts poor clinical outcomes in CRC patients

Overexpression of *SERPINE1* has been linked to adverse clinical outcomes in various cancer types^25,26^. Consistently, we found that the upregulation of *SERPINE1* was also associated with enhanced invasive disease and showed poorer clinical outcomes in multiple patient cohorts (**Supplementary Fig. S3B-G**). In this work we elucidate that not only *SERPINE1*, but high levels of RESIST-M1 (*SERPINE1*, *SMARCD3*) and low levels of RESIST-M2 (*CYP51A1*, *SC5D*, *ACAT2*, *HMGCR*, *HMGCS1*, *MVD*, *FDPS*) genes were significantly associated with CMS4 subtype of CRC that is associated with unfavorable clinical prognoses (**Fig. 5**).

**Figure 5.**
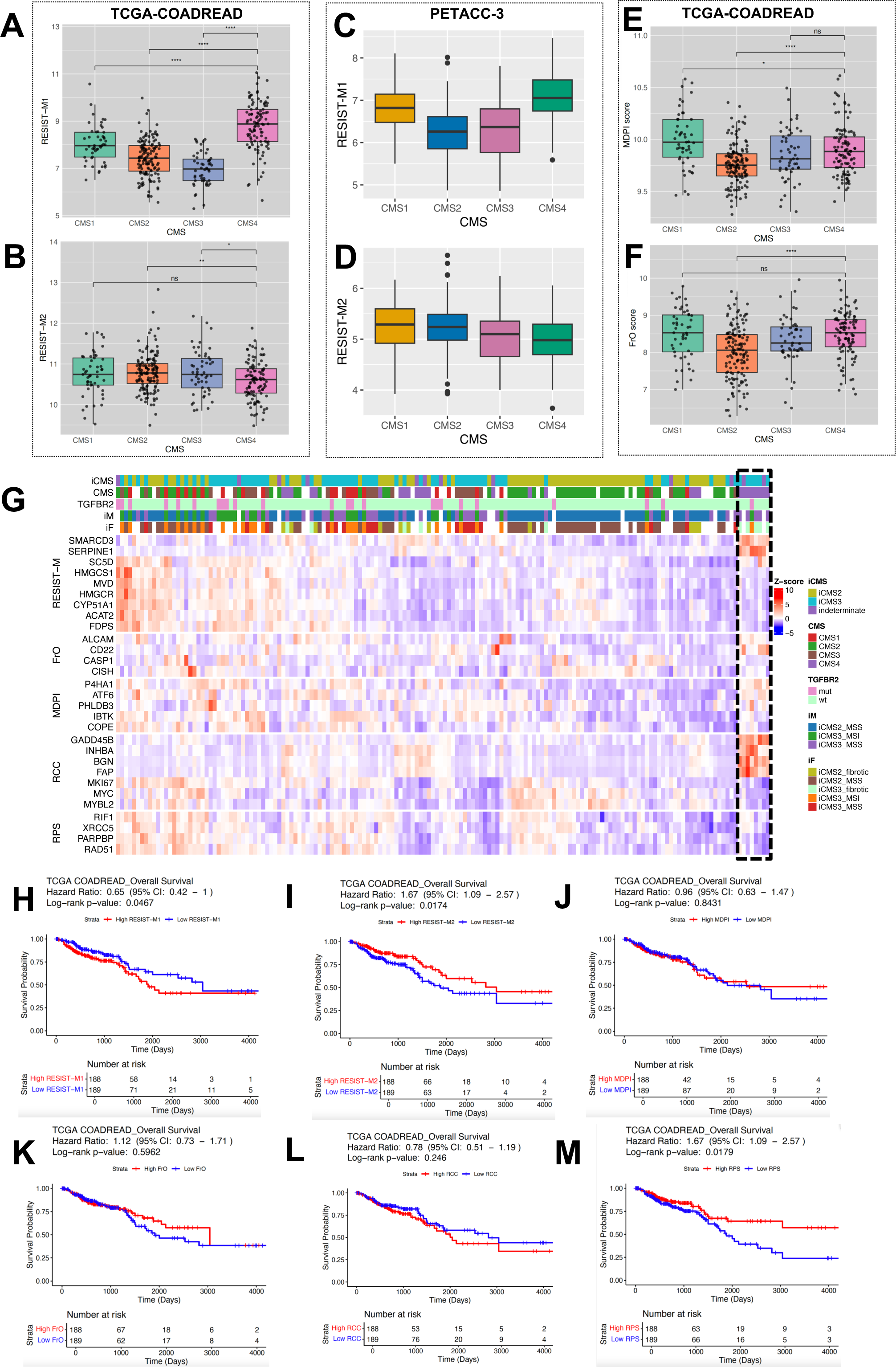
RESIST-M signature is significantly associated with CMS4 subtype and predicts poor clinical outcomes in CRC patients. **A-F, Boxplots showing average gene expression score for gene signatures across CMS subtypes in CRC patient datasets.** CMS4 has highest expression of RESIST-M1 genes and lowest expression of RESIST-M2 genes in TCGA-COADREAD (n= 377) (**A, B**) and PETACC-3 dataset (n = 604) (**C, D**). Mean of gene signatures reported for oxaliplatin resistance, such as MDPI (**E**) and FrO (**F**) are not unique to any CMS subtypes in TCGA-COADREAD data. RESIST-M1 gene score includes mean expression of *SERPINE1, SMARCD3*. RESIST-M2 gene score includes mean expression of *CYP51A1, SC5D, ACAT2, HMGCR, HMGCS1, MVD, FDPS*. MDPI gene score includes mean expression of *COPE, P4HA1, ATF6, IBTK, PHLDB3*. FrO gene score includes mean expression of *CD22, CASP1, CISH, ALCAM*. Statistical significance was determined using Wilcoxon rank-sum test where a p-value <0.05 is considered significant. **G, RESIST-M signature stratifies CMS4 and iCMS3 fibrotic patients.** Heatmap showing publicly available bulk RNA-seq data from Singapore colorectal cancer patients (n = 162, SG-BULK, ID: syn26720761) with RESIST-M signature. Patients with <u>high</u> expression of both *SERPINE1* and *SMARCD3* and <u>low</u> expression of *SC5D, FDPS, MVD, HMGCS1, HMGCR, CYP51A1, ACAT2* correlated predominantly with an CMS4-iCMS3-fibrotic subtype. iCMS3-fibrotic patients have been typically associated with poor relapse-free survival. In the heatmap, each column denotes data on a single CRC patient. Each row indicates the patient subtype based on available clinical data and z-scored values based on RESIST-M, FrO, MDPI, RCC, RPS gene expression from top to bottom. RCC gene score includes mean expression of twelve genes, out of which seven are used for analyses in this paper. The seven genes include stromal genes (*FAP, INHBA, BGN),* cell cycle genes *(MKI67, MYC, MYBL2)* and *GADD45B.* RPS gene score includes mean expression of four DNA repair genes-*RIF1, PARPBP, RAD51, XRCC5*. Clinical data shown in the legend includes patient subtype based on either iCMS, CMS, TGFBR2 mutation, iCMS-microsatellite (iM), or iCMS-fibrosis status (iF). **H-M, RESIST-M signature predicts poor clinical outcomes in CRC patients.** Kaplan Meier curves denoting overall survival for CRC patients in TCGA COADREAD dataset using RESIST-M1 (**H**), RESIST-M2 (**I**), MDPI (**J**), FrO (**K**), RCC (**L**), RPS (**M**) gene signatures.

Molecular characterization efforts, including Consensus Molecular Subtypes (CMS)^27^ and iCMS subtypes^28^, indicate limited benefit of oxaliplatin in CMS4 patients, a subtype which typically exhibits the poorest RFS and OS, along with the highest propensity for metastasis^5^. Similarly, iCMS3-fibrotic patients are associated with worst prognoses in metastatic CRC^28^. Remarkably, when we evaluated RESIST-M signature in two CRC datasets-a) TCGA COADREAD (n = 377, bulk RNA-seq of CRC patients with CMS annotations, **Fig. 5A-B**), b) PETACC-3 dataset^29^ (n = 604, bulk RNA-seq of CRC patients treated with 5FU-based chemotherapy, **Fig. 5C-D**), we found that RESIST-M1 and -M2 genes was most and least expressed, respectively, in CMS4 CRC patients compared to other molecular subtypes. This is in contrast with previously reported gene signatures on oxaliplatin resistance, labelled as MDPI^30^ and FrO^31^ signatures, which fail to differentiate between the different CMS subtypes (**Fig. 5E-F)**. This might be because the previously reported gene signatures on oxaliplatin resistance in CRC (MDPI, FrO) are derived from computational analysis of clinical datasets without a clear oxaliplatin-resistance induced metastatic phenotype. Additionally, we observed that RESIST-M genes (high expression of RESIST-M1 and low expression of RESIST-M2) could specifically stratify a subset of patients with only CMS4 and preferably iCMS3-fibrotic subtype (**Fig. 5G, Black dotted box**), whereas other gene signatures, such as MDPI^30^, FrO^31^, RCC^32^, RPS^33^ failed to identify the same set of patients in SG-BULK dataset (synapse ID: syn26720761). The RCC and RPS scores are frequently used in cancers to predict recurrence and mismatch repair respectively. The RCC score (also known as Oncotype DX test for colon cancer) includes a panel of 12 genes, with 7 cancer-related genes (stromal genes like *FAP, BGN, INHBA*, and cell cycle genes like *MKI67, MYC, MYBL2*, *GAD45B*) and 5 reference genes, and is used to predict CRC recurrence. However, the RCC score does not predict relative benefit from oxaliplatin ^32^. Intriguingly, we observe that patients with RESIST-M signature had high levels of stromal genes like *FAP, BGN, INHBA* and high *GAD45B*, while having low expression of cell cycle related genes like *MKI67, MYBL2, MYC*. Another reported marker of response to chemotherapy includes CDX2, wherein CRC patients lacking CDX2 expression are known to benefit from chemotherapy, but not oxaliplatin^34^. With our models, we provide evidence that oxaliplatin resistant cancer cells have low CDX2 expression; they display lower expression of both RPS and RCC genes, and are characterized by RESIST-M signature, which altogether probably make them unresponsive to adjuvant treatment with oxaliplatin (**Supplementary Fig. S3H-I**).

In accordance with the clinical outcomes linked to CMS4 CRC, RESIST-M gene signature could also predict poor prognoses in multiple datasets and performed better than previously reported gene signatures (**Fig 5H-M, Supplementary Fig. S4**). For instance, patients with high RESIST-M1 and low -M2 signatures were also associated with poor prognosis and outcome in TCGA COADREAD patients (**Fig 5H-I, Supplementary Fig. S4A)**, three publicly available patient cohorts (**Supplementary Fig. S4B-C)**, and in the PETACC-3 data which includes patients treated with adjuvant chemotherapy (**Supplementary Fig. S4D-F**). Particularly, low expression of RESIST-M2 genes was a strong predictor of poor relapse-free survival, specifically in CMS4 patients in the PETACC-3 cohort. Altogether, these data suggests that RESIST-M1 and M2 signature were anti-correlated and prognostic individually, but taken together, the RESIST-M signature can characterise pro-metastatic CMS4 CRC which can serve as a valuable tool to predict patient outcomes associated with oxaliplatin-resistance induced metastasis.

### Oxaliplatin-resistant HCT116 displayed enhanced therapeutic vulnerability against cholesterol and ROS-targeting drugs

The phenomenon of developing resistance to one drug often involves acquiring sensitivity to another^35^. In a quest to uncover such therapeutic vulnerabilities in our oxaliplatin-resistant models, we conducted an independent synthetic lethal drug screen using a panel of FDA-approved anticancer and kinase inhibitor libraries (**Fig. 6A-G** for HCT116-based models**, Fig. S5A-B** for SW480-based models). Notably, oxaliplatin-resistant HCT116 cells exhibited enhanced sensitivity to simvastatin, a cholesterol-lowering drug, both *in vitro* and *in vivo* (**Fig. 6A-G**). This discovery is consistent with our molecular findings, which indicate that oxaliplatin-resistant HCT116 cells downregulate cholesterol biosynthesis as an adaptive survival mechanism. Consequently, targeting cholesterol synthesis in these cholesterol-depleted cells seemed to improve the IC50 by atleast ∼2 folds compared to parental cell lines (**Fig. 6G)**. Previous reports provided evidence that depletion of cholesterol biogenesis can indeed be exploited to inhibit breast cancer metastasis^36^, however the use of statins has not yet been used to address oxaliplatin-induced metastasis, except limited reports on oxaliplatin-induced neurotoxicity^37^. Additionally, our screen identified elesclomol, a copper chelator known to induce oxidative stress and ferroptosis in CRC^38^, and inhibit CMS4 subtype that is associated with peritoneal metastasis^39^, as another promising candidate. The increased levels of reactive oxygen species (ROS) detected in oxaliplatin-resistant HCT116 cells (*data not shown*) as well as the CMS4-like metastatic properties of the resistant cell lines may underlie their heightened sensitivity to ROS-inducing agents. In summary, our findings uncover key therapeutic vulnerabilities in oxaliplatin-resistant CRC models, highlighting potential strategies to overcome resistance. These insights offer new avenues for targeting oxaliplatin-resistant tumors, including the pro-metastatic CMS4 subtype of colorectal cancer, especially those characterized by RESIST-M gene expression.

**Figure 6.**
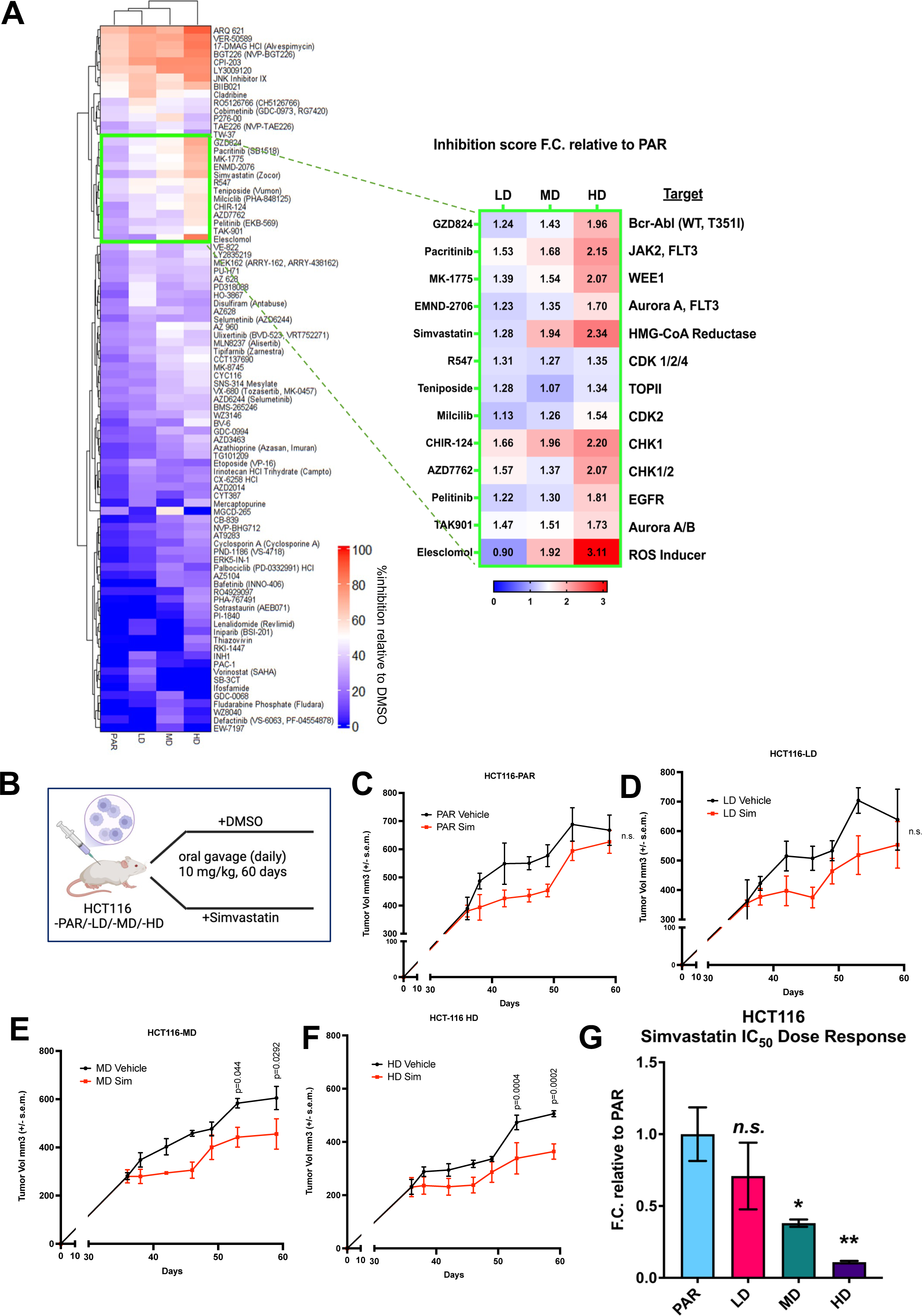
Simvastatin re-sensitizes resistant HCT116 to oxaliplatin treatment. **A, Drug screen assays on HCT116 oxaliplatin-resistant models.** Oxaliplatin resistant HCT116 models (HCT116 LD, MD and HD) were treated *in-vitro* with a library of FDA-approved drugs. Top hits identified are highlighted with their relative inhibition scores relative to PAR. Molecular targets associated with the identified molecules are mentioned alongside.e **B-G**, Validation of identified hit, simvastatin, on *in-vivo* tumor models. Scheme depicting generation of oxaliplatin resistant HCT116 models (HCT116 LD, MD and HD) and testing of simvastatin each of the models (**B**). Tumor growth curves of simvastatin treated HCT116 PAR (**C**), LD (**D**), MD (**E**) and HD (**F**) relative to drug vehicle treatment. Simvastatin IC_50_ fold change of HCT116 LD, MD and HD relative to PAR (**G**). Statistical significance was determined using Ordinary one-way ANOVA followed by Dunnett’s multiple comparisons test where a p-value <0.05 is considered significant.

## Discussion

Modelling chemoresistance and metastasis have largely relied on extended or supra-physiological drug exposure, which does not accurately reflect clinical treatment regimen^12,13^. In this study, we adopt a more clinically relevant, cyclical model of oxaliplatin administration with physiological dosing to generate drug-resistant CRC models. Using HCT116 and SW480 CRC cell lines, we administered clinically relevant doses of oxaliplatin, a commonly used first-line chemotherapy in CRC patients and evaluated the resulting cellular and molecular changes using a variety of experimental approaches, including bulk and single-cell transcriptomics, ELISA, Western blots, immunohistochemistry, and preclinical murine tumor models.

Our results demonstrate that mid to high doses of oxaliplatin induce both drug resistance and enhanced metastatic phenotypes, with differential resistant and metastatic potential observed between the two cell lines. Transcriptomic analysis revealed that drug resistance in both cell lines was associated with alterations in pro-inflammatory pathways such as JAK-STAT, NF-κB, and IFN-γ signaling, while tumor metastasis was more prominent in HCT116 compared to SW480. This disparity likely stems from intrinsic differences in their mutational landscapes, transcriptional profiles, and epigenomic states. Cancer cells with high migratory and metastatic potential often display mesenchymal features, while non-metastatic cells retain epithelial characteristics, such as polarity and cell-cell adhesion^40^. This is consistent with our findings, where oxaliplatin treatment led to upregulation of epithelial-to-mesenchymal transition (EMT)-associated pathways in HCT116, potentially explaining its dual resistant and pro-metastatic behavior.

Extensive molecular and phenotypic characterization of the resistant models revealed the upregulation of coagulation-related pathways and downregulation of metabolic pathways, including cholesterol biosynthesis, oxidative phosphorylation (OXPHOS), and fatty-acid metabolism. Through bulk and single-cell RNA sequencing we identified significant upregulation of *SERPINE1*, a gene involved in the coagulation pathway, alongside downregulation of cholesterol-related genes in oxaliplatin-resistant models. Based on these insights, we established the RESIST-M signature, comprising RESIST-M1 genes (e.g., *SERPINE1*, *SMARCD3*) and RESIST-M2 genes (e.g., *CYP51A1*, *SC5D*, *ACAT2*, *HMGCR*, *HMGCS1*, *MVD*, *FDPS*). Cells with positive RESIST-M signature (i.e. high RESIST-M1 and low RESIST-M2 expression), particularly in HCT116, exhibited both resistant and metastatic traits. Functional inhibition of *SERPINE1* using shRNA or pharmacological inhibitors like tiplaxtinin, resensitized resistant cells to oxaliplatin and reduced metastasis, establishing *SERPINE1* as a key player in oxaliplatin resistance-induced metastasis in CRC. Consistent with our findings, previous studies by independent groups have also demonstrated that *SERPINE1*, either tumor cell-intrinsic or cell-extrinsic, contributes to drug resistance and metastasis in various cancers, including, breast^19^ and head and neck cancers^18^. To the best of our knowledge, this study provides the first evidence implicating *SERPINE1* in driving oxaliplatin resistance and metastasis in CRC.

Mechanistically, our data suggest that oxaliplatin-resistant cells exploit both *SERPINE1* and cholesterol homeostasis for survival. Alterations in cellular cholesterol can disrupt the structure and composition of plasma membranes, particularly the caveolin-coated lipid rafts, which negatively regulate signaling pathways such as TGF-β^22^. Downregulation of cholesterol biosynthesis was linked to reduced membrane cholesterol levels in our data, impairing lipid raft function, and enhancing TGF-β signaling, which in turn upregulated *SERPINE1* and promoted EMT. This dysregulation in TGF-β receptor localization, due to decreased cholesterol biosynthesis, likely contributes to the sustained activation of the pathway in resistant cells, driving both resistance and metastasis in HCT116. These findings offer a unique strategy to target oxaliplatin-resistant cell lines, contrasting with cases of cancer where tumorigenesis is driven by increased cholesterol biogenesis and lipid metabolism^41^.

The clinical utility of this study is achieved by the validation of the RESIST-M signature in multiple independent CRC patient datasets that demonstrated its predictive value for poor prognosis. RESIST-M1 and RESIST-M2 gene signature were typically anti-correlated and characteristic of CMS4 CRC; RESIST-M1 genes were most and RESIST-M2 genes were least expressed in CMS4 CRC patients. Intriguingly, RESIST-M signature was particularly enriched in CRC patients with the CMS4 and iCMS3-fibrotic subtype in SG-BULK dataset. Indeed, these CRC subtypes are known to exhibit high metastatic potential, poor survival outcomes, and display minimal efficacy against oxaliplatin treatment, as evidenced by the NSABP C-07 and MOSAIC trials^3,4^. Finally, synthetic lethal drug screens revealed that RESIST-M positive oxaliplatin-resistant HCT116 models were particularly sensitive to simvastatin, a cholesterol-lowering drug, and elesclomol, an inducer of reactive oxygen species (ROS), known to enhance inhibit CMS4 CRC associated with peritoneal metastasis, via copper-dependent ferroptosis^38,39^. These findings suggest alternative therapeutic avenues for targeting resistant CRC, particularly the pro-metastatic CMS4 subtype, by combining cholesterol biosynthesis inhibitors or ROS inducers with conventional chemotherapies to overcome drug resistance and block metastatic progression.

In conclusion, our approach of generating clinically relevant drug-resistant CRC models against first-line chemotherapy allowed us to identify new mechanistic insights and actionable therapeutic targets that can be harnessed in the future to reverse therapy resistance and inhibit metastasis. In this work, we provide three critical resources: a) Improved oxaliplatin-resistant models, b) A novel gene signature to predict oxaliplatin resistance-induced metastasis, c) Therapeutic candidates targeting CMS-4/iCMS-fibrotic-like metastatic CRC cells (**Fig. 7)**. Our models represent aggressive CMS4/iCMS-fibrotic-like CRC profiles, facilitating further research in chemotherapy resistance and metastasis. The RESIST-M signature holds great promise for predicting oxaliplatin sensitivity in clinical settings, addressing a critical unmet need. As the RESIST-M signature is derived from resistant models with metastatic phenotypes, it is more prognostic for overall survival compared to previously reported gene signatures of oxaliplatin resistance, as validated in clinical cohorts. We identified promising therapeutic strategies, including *SERPINE1* inhibitors and statins, which demonstrated efficacy in reversing oxaliplatin resistance and metastasis in our models. We anticipate that future refinements of these approaches will improve treatment outcomes and provide valuable guidance for the effective administration of adjuvant chemotherapy in patients with oxaliplatin-resistant, pro-metastatic CRC.

**Figure 7.**
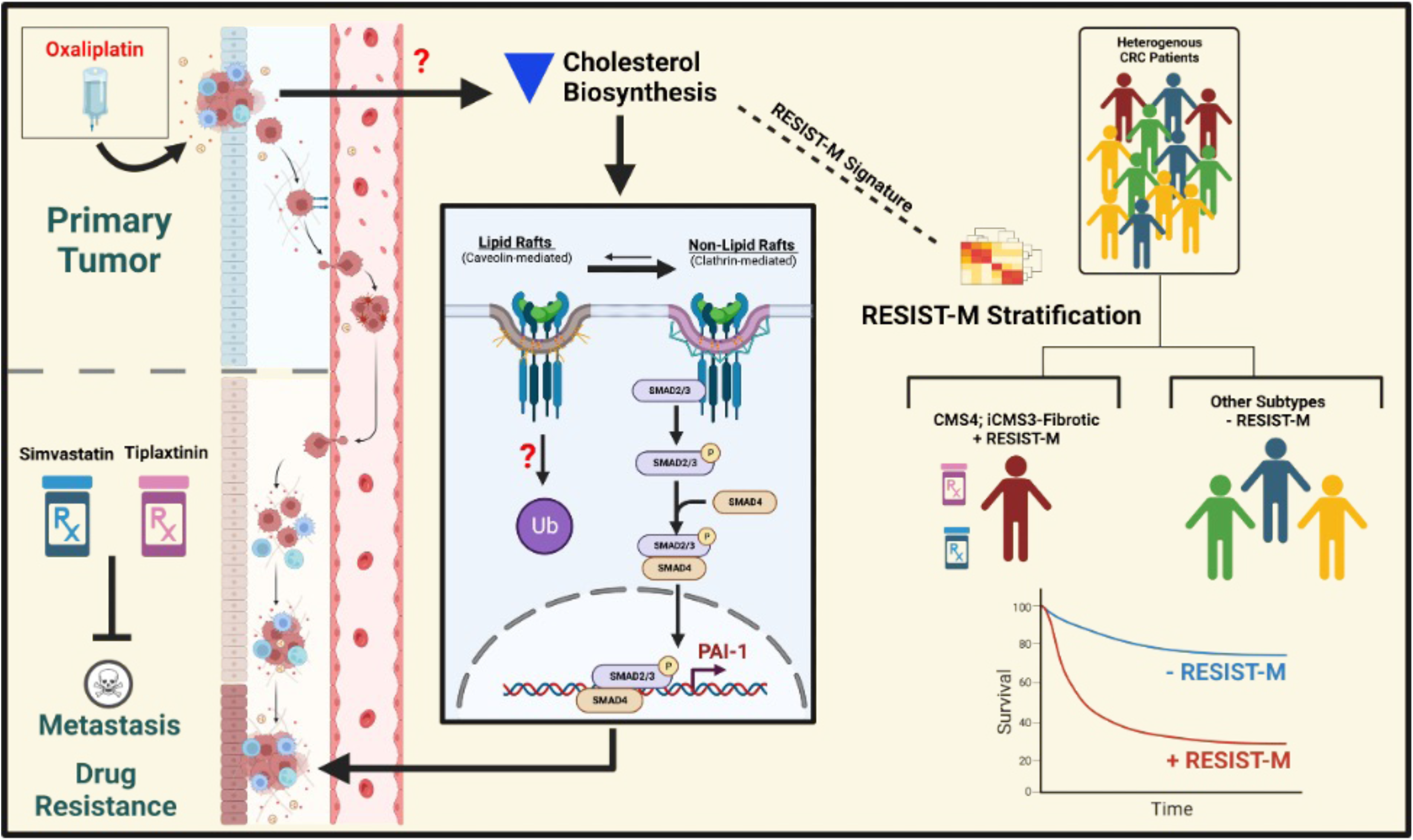
Schematic diagram depicting oxaliplatin-resistant models reveal gene signatures associated with, and strategies to overcome drug resistance-induced metastasis. Oxaliplatin-resistant CRC tumor cells reduce cholesterol biosynthesis that diminishes TGF-β receptor localization in caveolin-coated lipid rafts, thereby enhancing TGF-β signaling and activating expression of *SERPINE1*. This upregulated *SERPINE1* confers metastatic and invasive properties to the resistant CRC tumor cells. A novel *SERPINE1*-based transcriptomic signature, RESIST-M, can stratify CRC patients into iCMS3-CMS4 subtype, that are associated with poor survival outcome, metastatic relapse, and lack of response to oxaliplatin in patients. Inhibition of *SERPINE1* or cellular cholesterol levels using statins diminishes metastatic colonization, and re-sensitize resistant tumor cells to oxaliplatin.

## Acknowledgements

This work was supported by Singapore National Medical Research Council’s grants (NMRC/CIRG/1439/2015) awarded to R.D, IBT; NMRC/OFIRG/0025/2019 to RD; National Research Foundation grant NRF-CRP26-2021-0005 to RD; the Industry Alignment Fund Pre-Positioning Program’s fund (IAF-PP Cat 1 grant to RD supporting a joint lab with PerkinElmer) and the A-STAR GIS-Core research support.

## Materials and Methods

### Cell culture

All cell lines were grown in standard cell culture conditions (37°C, 5% CO2). HCT116 and SW480 were obtained from ATCC and maintained in Modified McCoy’s 5A media containing L-Glutamine (Gibco) supplemented with 10% FBS, 1% penicillin/streptomycin and 1% sodium pyruvate. All cell cultures were routinely tested for the presence of mycoplasma using the MycoAlert Mycoplasma Detection Kit (Lonza, USA) following manufacturer’s protocol.

### Generation of oxaliplatin resistant cell lines

Oxaliplatin (CAS#: 61825-94-3, Tocris) was dissolved in DMSO. Cells were treated for 72 hours with different concentrations of oxaliplatin-0.5 µM, 5 µM and 80 µM, before changing to drug-free media. The drug-free media was replaced every alternate day until cells became confluent. Cells are sub-cultured once they reached confluency and treated at the same dose for 72 hours.

This treatment cycle was repeated for a total of 10 times. The resulting oxaliplatin resistant cell lines were then used in subsequent experiments and routinely maintained in drug-free media.

### In vitro dose-response assay

Approximately 5000 – 8000 cells were seeded into each well of a 96-well plate and incubated for 48 hours. Cells were then treated with oxaliplatin for 72 hours at nine different doses serially diluted 3-fold with a starting dose of 1000 µM. DMSO was used as a no-drug control. Cell viability was measured using CellTiter-Glo (Promega) following manufacturer’s protocol. Data points were plotted in Prism 9 and analyzed using the nonlinear regression function “log(inhibitor) vs. normalized response - Variable slope”.

### Cell proliferation assays

Real-time proliferation kinetics was done using IncuCyte (Essen Bioscience) following manufacturer’s protocol. Briefly, 5000 cells were seeded and cultured in 96-well plates, and images were taken every hour for the next 72 hours. Percentage cell confluence was quantified using the in-built IncuCyte software.

### Transwell invasion and migration assays

Cultrex 96 Well BME Cell Invasion Assay kits (RnD Systems) were used for transwell invasion and migration assays following manufacturer’s protocol. Briefly, cells seeded prior to assay and grown to about 70% confluency were treated with 10 ug/mL of Mitomycin C (CAS#: 50-07-7, MedKoo Biosciences) for 2 hours. 5 x 10^5^ cells resuspended in serum free media were seeded into each upper chamber well in triplicates. Wells used for invasion assays were pre-coated overnight with 0.1X Matrigel (Corning). Uncoated wells were used for migration assays. Complete media was added to the lower chamber. Percentage invasion was determined by measuring the number of cells that had invaded to the bottom chamber divided by total number of cells seeded.

### RNA sequencing & Gene Set Enrichment Analyses

RNA was extracted using the RNeasy Plus Mini Kit (QIAGEN) following manufacturer’s protocol. Yield and integrity of RNA were determined using the Agilent 2100 Bioanalyzer (Agilent Technologies) following manufacturer’s protocol. Sample preparation for the Bioanalyzer was done using the Agilent RNA 6000 Nano Kit (Agilent Technologies, USA) following manufacturer’s protocol. RNA sequencing was done at the Next Generation Sequencing Platform in the Genome Institute of Singapore, A*STAR. All pre and post processing steps, including bioinformatics analyses were performed by the sequencing platform. A list of read counts was provided as the final output of the sequencing. RNA sequencing data was further analyzed on various bioinformatics platforms. In general, differentially expressed genes were obtained using the DESeq2 package^42^. Subsequently, GSEA using hallmark gene sets from the Molecular Signatures Database (MSigDB)^43,44^ was done using the fgsea package^45^. All packages and pipelines were processed using R on RStudio

### Single-cell RNA sequencing analyses

In-house generated single-cell datasets from HCT116 PAR/LD/MD/HD cell lines were processed through the Cellranger 6.0.1 pipeline, aligned to the human reference genome (GrCh38). The raw feature-barcode matrix was loaded into Seurat (4.3.0). Cells of low quality (fewer than 200 genes per cell) and cells expressing rare genes (those expressed in fewer than 30 cells) were excluded from the analysis. Cells with over 20% mitochondrial gene content were also removed. Potential cell doublets were filtered out by first setting an upper limit of 6000 genes detected per cell and then consequently by using the DoubletFinderpackage. The data was normalized with ScaleData() using a scaling factor of 10000, and gene expression was scaled to unit variance and a mean of 0. Principal component analysis was performed using the RunPCA function for dimensionality reduction, with an elbow plot determining the inflection point where variance changes became insignificant. The neighborhood graph for clustering was calculated using FindNeighbors, and cell clustering was done with the FindClusters function using louvain clustering. UMAP of the clusters were made using RunUMAP() function and a splitUMAP() function was used to compare the differences in clusters across different conditions. CellPropPlot() was used to assess the proportion of cell clusters across different conditions. Signature scores were calculated using the UCell package. Lastly, the expression of Resist-M signatures was verified across the different LD/MD/HD conditions using DotPlot() or VlnPlt() function.

### Quantitative RT-PCR

RNA was extracted as per previously described. Nanadrop was used to determine the concentration and quality. cDNA was synthesized using SuperScript IV VILO Master Mix (11756500, Invitrogen) following manufacturer’s protocol. qPCR was done using the KAPA SYBR FAST qPCR Master Mix (2X) Kit (KAPA BIOSYSTEMS) following manufacturer’s protocol and qPCR cycles were done on QuantStudio 7 Flex Real-Time PCR System (Applied Biosystems). Primers were purchased from Integrated DNA Technologies, Singapore. Primer sequences are found in table 1. All qPCR runs were done with technical quadruplicates and three independent biological replicates.

### ELISA

Measurement of PAI-1 (BMS2033, Thermo) and TGF-β1 (ab100647, abcam) levels was performed following manufacturer’s protocol. A suitable number of cells were first seeded in 6-well plates supplemented with complete media. 24 to 48 hours later, spent media was replaced with serum free media and incubated for a further 24 to 48 hours. Supernatant was collected for downstream ELISA assays. Cells were lysed in RIPA and subjected to protein quantification using BCA assay for normalization.

### Western Blot

Western blot was done as per standard protocol. Briefly, cells were lysed in RIPA buffer (89901, Thermo) supplemented with Protease Cocktail Inhibitor (4693116001, Sigma) and PhosSTOP (4906837001, Sigma). Lysates were cleared by centrifugation and subjected to protein quantification using the Pierce BCA Protein Assay Kit (23225, Thermo) following manufacturer’s protocol. NuPAGE LDS sample buffer (NP0008, Invitrogen) supplemented with NuPAGE reducing agent (NP0009, Invitrogen) was added to the protein lysates and subjected to SDS-PAGE. Wet transfer was done onto PVDF membrane (IPFL85R, Merck) and blocked for 1 hour with TBS blocking buffer (927-60001, LI-COR) or 1.5% non-fat milk (#1706404, Biorad), and subsequently incubated with primary antibodies overnight at 4°C. NIR imaging was done using the LI-COR Odyssey CLx system and Image Studio Ver 4.0. Chemiluminescence imaging was done using the Azure 600 (Azure Biosystems) after incubation with HRP substrate (K-12042-D20, Advansta). List of antibodies and dilutions used are found in table 2.

### Animals

6 to 8 weeks old male NSG (NOD.Cg-Prkdc^scid^ IL2rg^tm1Wjl^/SzJ) immunocompromised mice purchased from Invivos Pte Ltd (Singapore) were used in all animal studies. Animals were group housed in individually ventilated cages in the Biological Resource Centre, A*STAR, Department 3. Room lighting was set to a 12-hours light-dark cycle as recommended by the National Advisory Committee for Laboratory Animal Research (NACLAR). Animals were provided with irradiated Altromin 1324 diet and autoclaved water, *ad libitum*. The protocol was approved by the Biological Resource Centre Animal Use and Care Committee (IACUC #181372), and animals were maintained in accordance with guidelines from the American Association of Laboratory Animal Care (AALAC). No animals were excluded from any studies unless specified. Sex is not a biological variable for our studies.

### In vivo studies

#### Tumor growth kinetics and spontaneous metastasis

Each mouse was injected subcutaneously in the left flank with 0.5 to 1 million cells resuspended in serum free McCoy’s media and 50% Matrigel (354234, Corning). Tumor growth was tracked weekly by measuring using Vernier calipers. Tumor volume was determined by (L x W^2^)/2, where L is the longest axis and W is the perpendicular axis. Eight weeks post inoculation, resection of the primary tumor was done under anesthesia and mice were left to recover with proper post-operative care. Lungs were then harvested to assess for metastasis three to four weeks after surgery. Lungs were first perfused with 10% neutral buffered formalin (HT501128, Sigma) during harvesting and fixed in the same fixative for at least 48 hours. Paraffin blocking, sectioning and relevant H&E staining were done at the Advanced Molecular Pathology Laboratory (AMPL) in the Institute of Molecular & Cell Biology (IMCB), A*STAR. Tumor measurements were not blinded.

#### In vivo effects of oxaliplatin treatment on spontaneous metastasis

Mice were injected with treatment naïve HCT116 or SW480 cells as described previously. Clinical grade oxaliplatin was kindly provided for by Dr Clarinda Chua from the National Cancer Center of Singapore. Drug was diluted with water for injection (B.Braun) to a final concentration of 1 mg/mL before treatment. Mice were randomized when the average cohort tumor size reached 200 mm^3^ to 300 mm^3^. Treatment groups received 5 mg/kg of oxaliplatin weekly, *i.p.*, for 5 consecutive weeks, while vehicle group was injected with water for injection at 0.1 ml per 10 g of body weight.

Tumor resection was done one-week after the final dose of treatment, and animals were sacrificed three to four weeks after surgery for organ harvesting as per previously described. Drug treatments and tumor measurements were not blinded.

#### Impact of concomitant administration of tiplaxtinin and oxaliplatin on metastasis

Mice were injected with naïve or parental HCT116 cells and randomized as described previously. Tiplaxtinin (CAS # 393105-53-8, Tocris or Selleckchem) was dissolved in DMSO to a concentration of 50 mg/mL, and then added to corn oil (Sigma) to a final concentration of 5 mg/mL. Treatment group received 20 mg/kg of tiplaxtinin daily (Monday to Friday), *p.o.*, for five consecutive weeks. Vehicle groups received only corn oil added with equal amounts of DMSO at 0.1ml per 10g of body weight. All mice for this experiment received Oxaliplatin at 5 mg/kg once a week, *i.p.*, for five consecutive weeks. Tiplaxtinin treatment began with the first dose of oxaliplatin. Tumor resection was done one-week after the final dose of treatment, and animals were sacrificed three to four weeks after surgery for organ harvesting as per previously described. Drug treatments and tumor measurements were not blinded.

#### In vivo effects of simvastatin treatment on oxaliplatin resistant tumor models

Mice were injected subcutaneously with either parental HCT116 (HCT116-PAR) or the oxaliplatin resistant cell line models (HCT116-LD, MD, HD). Once the tumor size reached approximately 200 mm^3^ to 300 mm^3^ mice were randomized into either control or drug treatment group for each cell line as described previously. Control and drug treatment group received DMSO and simvastatin respectively. Simvastatin (CAS # 79902-63-9, Tocris) was dissolved in DMSO to a concentration of 50 mg/mL, and then added to corn oil (Sigma) to a final concentration of 2 mg/mL. Drug treatment group received 10 mg/kg of simvastatin daily (Monday to Friday) by oral gavage. Drug treatments and tumor measurements were not blinded.

### Slide scan acquisitions and image analyses

Tissue slides were scanned using the Vectra Polaris Automated Quantitative Pathology Imaging System (Akoya Bioscience) and analyzed using both inForm Tissue Finder version 2.4 and Phenochart version 1.0 software by Akoya Bioscience. Slides were scanned at 20X magnification and exported as TIFFs to be analyzed with InForm.

To determine the severity of pulmonary metastasis, image analysis of tumor nodules and normal lung tissue areas were quantified using the threshold algorithm in InForm. Metastasis score is calculated using the following formula: (Area of all nodules/ Total area of tissue) x 100.

### Genetic knockdown using shRNA

To generate stable knockdown lines, suitable bacteria clones from the RNAi Consortium (TRC) shRNA Library (Broad Institute) were selected and sub-cultured in terrific Broth (CUS-4051-1L, Axil Scientific) supplemented with 1X carbenicillin (10177012, ThermoFisher). Plasmids were extracted using the Plasmid Plus Midi kit (12945, QIAGEN) or Plasmid Plus Maxi kit (12963, QIGEN) following manufacturer’s protocol. Purity was determined using the Nanodrop 8000.

Custom shRNAs against PAI-1 were designed using the BLOCK-iT™ RNAi Designer (ThermoFisher) and cloned into the pLKO.1 vector. The TRC cloning vector was a gift from David Root (Addgene plasmid #10878; RRID: Addgene_10878). Lentivirus was generated in HEK293T cells using Lipofectamine 3000 Transfection kit (L3000150, Thermo) according to manufacturer’s protocol. A total of 9 ug of shRNA plasmid and 3 ug each of the third-generation lentiviral packaging plasmids were used for the transfection mix. The three lentiviral plasmids, pMD2.G (Addgene plasmid #12259; RRID: Addgene_12259), pMDLg/pRRE (Addgene plasmid #12251; RRID: Addgene_12251), and pRSV-Rev (Addgene plasmid #12253; RRID: Addgene_122-53) were gifts from Didier Trono. Transfection lasted for about 16 to 24 hours before a change to appropriate, fresh media. After 24 hours, supernatant containing the viral particles was collected and centrifuged at 1,000 rpm for 5 minutes and filtered using a 0.45um filter before adding onto target cells supplemented with 1X polybrene (sc-134220, Santa Cruz). Cells were transduced for 24 to 48 hours. Puromycin selection was then done for at least two passages to establish stably transduced cell lines. Knockdown of target genes was validated with qPCR, western blot, and / or ELISA. Sequences of control and PAI-1 shRNAs used are listed in Table 3.

### Immunofluorescence staining

Cells were seeded and cultured in standard cell culture conditions for 24 hours prior to staining on 8-well imaging slides (80826, Ibidi). Lipid raft staining was done using the lipid raft staining kit (V34404, ThermoFisher) according to manufacturer’s protocol. Cells were then subsequently co-stained with primary antibody using TGFbRII (79424S, CST) and secondary antibody using AF647 (A-21244, Thermo). Images were taken using the Nikon Ti-E TIRF System.

### Lipid rafts sucrose density fractionation

Lipid raft fractionation protocol was adapted from Zuo et. al^46^. Briefly, cells were seeded and grown to near confluence in four 100-mm dishes. During harvesting, cells were washed twice with ice-cold PBS and scraped with 0.75 mL of 500 mM sodium carbonate, pH 11.0 and incubated on ice for 10 minutes. Mechanical homogenization was done using 10 strokes of a Dounce homogenizer followed by three 20-seconds bursts of sonication on ice. Homogenates were adjusted to 42.5% sucrose by adding equal volumes of 85% sucrose in 2X HBS (25 mM HEPES, pH6.5, 150 mM NaCl) in an ultracentrifuge tube (344060, Beckman Coulter). 3.5 mL of 35% and 5% sucrose were overlayed to create a discontinuous sucrose gradient. Samples were subjected to ultracentrifugation of 260,000 x g for 16 hours at 4°C using the Optima XPN-80 and SW40Ti rotor (Beckman Coulter). Twelve 1-mL fractions were collected from the top and subjected to immunoblotting.

### Patient cohorts and survival analyses

#### Datasets

Normalised bulk RNA-sequencing data for TCGA-COADREAD, TCGA-COAD and -READ datasets were obtained from UCSC Xena browser. Normalised counts for bulk RNA-sequencing data were obtained for PETACC-3 dataset as described^29^ and SG-BULK dataset (synID: syn26720761). Robust Multichip Average/Frozen robust multiarray analysis (RMA/FRMA) normalized microarray datasets^27^ (GSE14333, GSE37892, GSE39582) were obtained from Synapse ID: syn2634742.

#### Analyses of transcriptomic data from public datasets

Boxplots for gene signatures across CMS subtypes is shown for TCGA-COADREAD data (n = 377, with CMS annotations) and PETACC-3 data (n = 604). CMS annotations for patients in TCGA-COADREAD data was extracted from Colorectal Cancer Subtyping Consortium (CRCSC) available at Synapse ID: syn2623706. The CMS was calculated with the reference CMS classifier (https://github.com/Sage-Bionetworks/CMSclassifier) for PETACC-3 dataset. Gene signatures shown include RESIST-M1(*SERPINE1, SMARCD3*), RESIST-M2 (*SC5D, FDPS, MVD, HMGCS1, HMGCR, CYP51A1, ACAT2*), MDPI^30^ (COPE, P4HA1, ATF6, IBTK, PHLDB3), FrO^31^ (*CD22, CASP1, CISH, ALCAM*).

Heatmap comparing RESIST-M genes to reported gene signatures of oxaliplatin resistance (MDPI^30^, FrO^31^), CRC recurrence (RCC^32^) and DNA repair gene signature (RPS^33^) was done using publicly deposited bulk RNA-seq data from Singapore colorectal cancer patients (n = 162, SG-BULK, Project SynID: syn26720761^28^). Genes were tabulated using the available clinical annotations for iCMS, CMS, TGFBR2 mutation, iCMS-microsatellite (iM), or iCMS-fibrosis status (iF). Salmon TPM counts of RESIST-M signature genes (*SERPINE1, SMARCD3*, *SC5D, FDPS, MVD, HMGCS1, HMGCR, CYP51A1, ACAT2*) and rest of the signatures were z-scored using scale() function in R and heatmap was made using heatmap() function using R version 4.2.2. Patients with <u>high</u> expression of both *SERPINE1* and *SMARCD3*, along with <u>low</u> expression of *SC5D, FDPS, MVD, HMGCS1, HMGCR, CYP51A1, ACAT2* correlated predominantly with an CMS4-iCMS3-MSS-fibrotic subtype. RCC gene score includes expression of twelve genes, out of which seven are used for analyses in this paper. The seven genes include stromal genes (*FAP, INHBA, BGN*), cell cycle genes (*MKI67, MYC, MYBL2*) and GADD45B^32^. RPS gene score includes expression of four DNA repair genes-RIF1, PARPBP, RAD51, XRCC5^33^.

Kaplan-Meier (KM) survival curves were generated to evaluate the prognostic significance of gene signatures: RESIST-M1, RESIST-M2, MDPI, FrO, RCC and RPS using TCGA-COADREAD dataset. Overall survival (OS) of patients were stratified into high- and low-expression groups based on median expression levels. Survival analysis was performed using the log-rank test, and hazard ratios were calculated using the Cox proportional hazards model.

KM curves plotted for RESIST-M1 and -M2 signatures together is represented by median of all signature genes expression (9 genes: *ACAT2, CYP51A1, FDPS, HMGCR, HMGCS1, MVD, SC5D, SERPINE1, SMARCD3*) in the sample. For gene *SMARCD3* & *SERPINE1*, a reciprocal is done with the following formula, before combining with other genes to get the median expression: min + (max - min)*(max - expression)/(max - min). Each sample will be assigned a high (1) or low (0) status depending on if its expression is higher or lower than the median signature gene expression threshold. Median signature gene expression threshold is obtained from median of all samples signature expression. Kaplan-Meier survival plot is using R "ggsurvplot" and "survfit" function with log-log confidence interval type.

Gene expression data were obtained from the PETACC-3 trial as previously described^29^. The M1 and M2 scores were obtained by calculating the mean expression of M1 and M2 genes for each patient. The score was considered "high" if above the median, "low" otherwise. Survival was modelled with Cox regression and the log-rank test. We used the survminer package to plot Kaplan-Meier curves. The CMS was calculated with the reference CMS classifier (https://github.com/Sage-Bionetworks/CMSclassifier) for PETACC-3 dataset. P-values were considered significant if below 0.05.

### Synthetic lethal drug screen

About 3000 cells were seeded in each well of black 384-well plates and incubated under standard cell culture conditions for 48 hours. Culture media was then refreshed in the morning, and drugs from the Anti-Cancer (L3000, Selleck Chemicals, USA) and Kinase Inhibitor (L1100, Selleck Chemicals, ISA) compound library were added at a concentration of 1 µM to each well in the afternoon. Cells were treated for 72 hours, and cell viability was determined using CellTiter-Glo (Promega) following manufacturer’s protocol. Data obtained was analyzed using EARO High-throughput Phenomics Database Portal. Percentage activity was calculated luminescence value of drug treated wells over DMSO control.

### Statistical Analyses

All experiments were performed in at least three biological replicates, and data for descriptive statistics are expressed as standard error of mean (s.e.m.) unless otherwise stated. Methods of statistical analysis are described in text. A p-value of less than 0.05 is considered statistically significant. All analyses were done with GraphPad PRISM v9 and v10.

---

**Supplementary Figure 1. Characterisation of SW480 oxaliplatin resistant model**

**A,** Representative dose response curves of SW480 parental (PAR), low dose (LD), mid dose (MD) and high dose (HD) to 48 hrs of oxaliplatin treatment.

**B,** Oxaliplatin IC_50_ fold change of SW480 LD, MD and HD relative to PAR. Statistical significance was determined using Ordinary one-way ANOVA followed by Dunnett’s multiple comparisons test where a p-value <0.05 is considered significant.

**C,** 4x phase contrast images of cell morphologies.

**D,** Transwell invasion assay in SW480 PAR, LD, MD and HD. Statistical significance was determined using Ordinary one-way ANOVA followed by Dunnett’s multiple comparisons test where a p-value <0.05 is considered significant.

**E,** Transwell migration assay in SW480 PAR, LD, MD and HD. Statistical significance was determined using Ordinary one-way ANOVA followed by Dunnett’s multiple comparisons test where a p-value <0.05 is considered significant.

**F,** Representative H&E staining of lung sections for **s**pontaneous lung metastasis in mice harboring SW480 PAR, LD, MD and HD xenografts.

**G,** Quantification of nodule positive area vs total lung area in mice harboring SW480 PAR, LD, MD and HD xenografts (n=5). Statistical significance was determined using Ordinary one-way ANOVA followed by Dunnett’s multiple comparisons test where a p-value <0.05 is considered significant.

**H,** Cell proliferation rate of HCT116 tracked using Incucyte.

**I,** Cell proliferation rate of SW480 tracked using Incucyte.

**J,** Tumor growth kinetics *in vivo* of SW480 PAR, LD, MD and HD xenografts (n= 5). Statistical significance was determined using Ordinary 2way ANOVA followed by Dunnett’s multiple comparisons test where a p-value < 0.05 is considered significant.

**K,** Corresponding images of tumor of SW480 PAR, LD, MD and HD xenograft at day 40.

**L,** Three representative H&E staining of lung sections from mice harboring SW480 PAR xenografts treated with vehicle or oxaliplatin for spontaneous metastasis.

**M,** Quantification of nodule positive area vs total lung area in mice harboring SW480 PAR, LD, MD and HD xenografts (n=4). Statistical significances was determined using two-tailed unpaired t-test. A p-value of <0.05 is considered significant.

**Supplementary Figure 2. Hallmarks of the resistant models using bulk and single-cell RNA seq analyses reveal resistant cells to be terminally differentiated**

**A,** Gene Set Enrichment Analysis (GSEA) using Differentially Expressed Genes (DEGs) from bulk RNA-seq analyses of HCT116 LD/ MD/ HD vs PAR. Red and blue indicates a positive and negative Normalized Enrichment Score (NES), respectively. All hallmarks shown here are significantly enriched.

**B,** Gene Set Enrichment Analysis (GSEA) using Differentially Expressed Genes (DEGs) f from bulk RNA-seq analyses of SW480 LD/ MD/ HD vs PAR. Red and blue indicates a positive and negative Normalized Enrichment Score (NES), respectively. All hallmarks shown here are significantly enriched.

**C,** Dot plot showing differentially expressed genes (DEGs) of the 5 identified clusters (cl 0-4)

**D,** Violin plot showing *SERPINE1* expression across different cell states labelled by annotations (cell clusters cl 0-4)

**E,** Violin plot showing *SERPINE1* expression across samples: HCT116 LD, MD, HD and PAR. **F,** UMAP projection and sample type annotation of scRNA-seq data from the four scRNA-seq libraries: HCT116-PAR/LD/MD/HD

**G,** UMAP projection showing corresponding Cytotrace score

**H,** Violin plot showing cytotrace score for each sample type

**I,** Trajectory analysis including all cell clusters, except cl 3, using Monocle 2 with corresponding annotations for sample type

**J,** Trajectory analysis including all cell clusters, except cl 3, using Monocle 2 with corresponding annotations for pseudotime

**Supplementary Figure 3. *SERPINE1* expression correlates with enhanced metastatic properties in human CRC patients and analyses of reported gene scores in our models A**, Scheme showing genes encoding for enzymes involved in the cholesterol mevalonate biosynthesis pathways.

**B,** *SERPINE1* expression in patients from the COAD TCGA dataset with and without perineural invasion.

**C,** *SERPINE1* expression in patients from the COAD TCGA dataset with and without LN metastasis.

**D,** *SERPINE1* expression in patients from the COAD TCGA dataset with and without venous invasion.

**E,** *SERPINE1* expression in patients from the COAD TCGA dataset with and without lymphatic invasion.

**F,** Overall Survival (OS) of patients from the TCGA COAD dataset stratified by *SERPINE1*.

**G,** Progression Free Survival (PFS) of patients from three publicly available datasets stratified by

*SERPINE1*.

**H,** Violin plot showing expression of CDX2 in our oxaliplatin resistant models.

**I,** Dot plot showing expression of RPS and selected RCC genes (left) and RESIST-M genes (right) in our oxaliplatin resistant models. Labels denote HCT116 parental (PAR) and dose of oxaliplatin used to generate the models, i.e. low dose (LD), mid dose (MD) and high dose (HD).

Statistical significance was determined using Ordinary one-way ANOVA followed by Dunnett’s multiple comparisons test where a p-value <0.05 is considered significant.

**Supplementary Figure 4. RESIST-M expression correlates with poor prognosis in human CRC patients**

**A,** Kaplan Meier curves denoting Disease Specific Survival (DSS) of patients from the TCGA COAD dataset stratified using RESIST-M signature. “+RESIST-M” refers to combination of high RESIST-M1 genes (i.e. *SERPINE1* and *SMARCD3)* and low RESIST-M2 genes (i.e. *SC5D, FDPS, MVD, HMGCS1, HMGCR, CYP51A1, ACAT2* ).

“-RESIST-M” refers to the opposite combination i.e. low M1 and high M2.

**B,** Relapse-free survival (RFS) of patients from publicly available datasets GSE14333, 37892, 39582 stratified using RESIST-M signature.

**C**, RFS of patients from publicly available datasets GSEA14333, 37892, 39582 stratified using each gene in the RESIST-M signature. Red and blue curves denote high and low expression of each gene respectively.

**D-F**, RFS for CRC patients in PETACC-3 dataset using RESIST-M2 gene signature for CMS4 alone (**D**), CMS1-3 (**E**), all CMS (**F**) subtypes. Red and blue curves denote low and high expression of each gene respectively.

Statistical significance was determined using Ordinary one-way ANOVA followed by Dunnett’s multiple comparisons test where a p-value <0.05 is considered significant.

**Supplementary Figure 5. Therapeutic targeting of oxaliplatin resistant SW480 models**

**A,** Synthetic lethal drug screen using the Anti-Cancer and Kinase Inhibitor library in SW480 resistant models

**B,** Identified interventional strategies for SW480 resistant models. Inhibition scores of SW480 LD, MD and HD relative to PAR using available drug molecules. The scores denote fold changes in IC_50_ compared to parental cell lines.

Statistical significance was determined using Ordinary one-way ANOVA followed by Dunnett’s multiple comparisons test where a p-value <0.05 is considered significant.

## Notes

### Competing Interest Statement

The authors have declared no competing interest.

