## Supplementary figures and tables for "Modelling oxaliplatin resistance in colorectal cancer reveals a *SERPINE1*-based gene signature (RESIST-M) and therapeutic strategies for pro-metastatic CMS4 subtype"

### Supplementary Figure 1. Characterisation of SW480 oxaliplatin resistant model

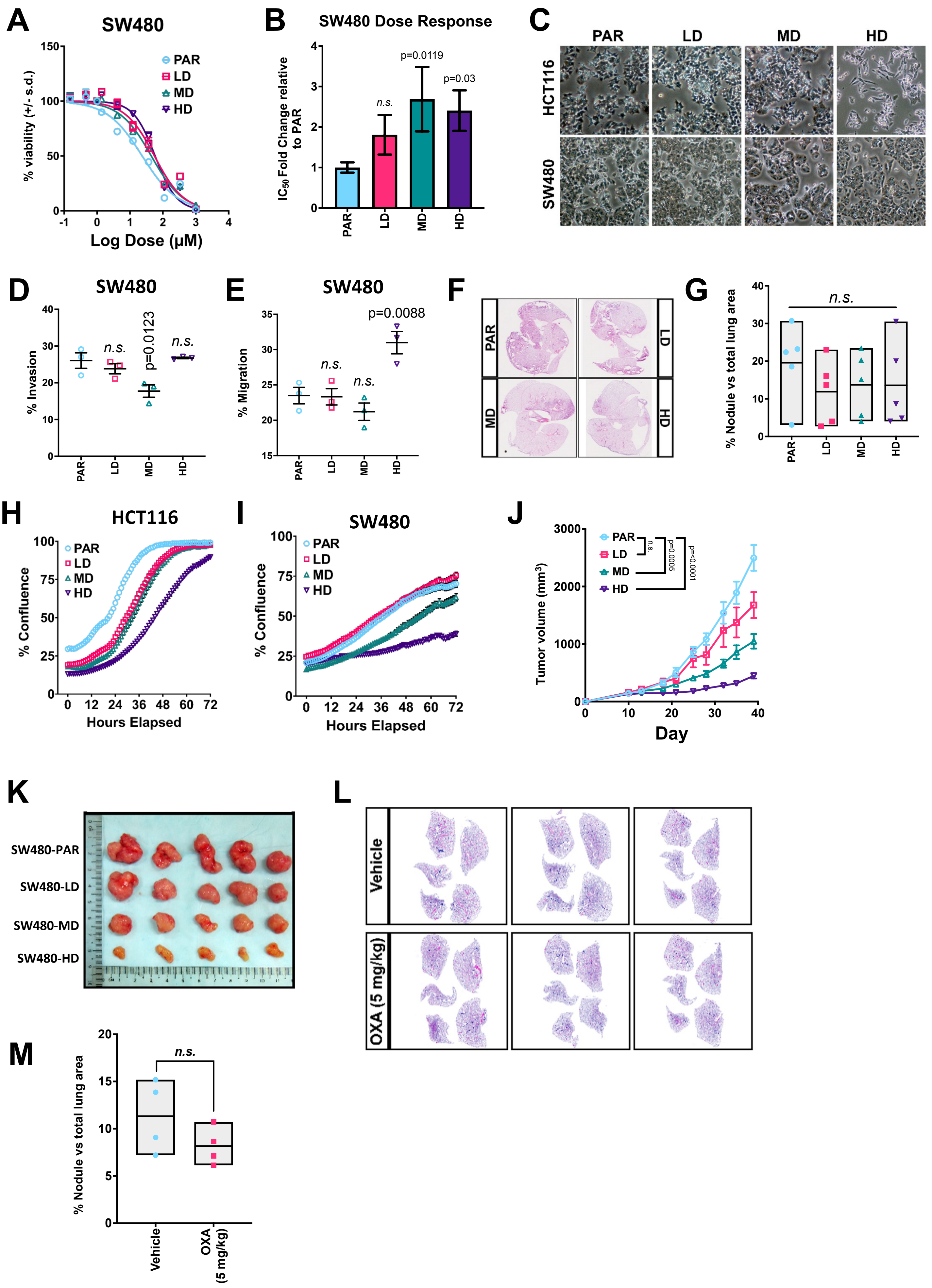

**Supplementary Figure 2. Hallmarks of the resistant models using bulk and single-cell RNA seq analyses reveal resistant cells to be terminally differentiated**

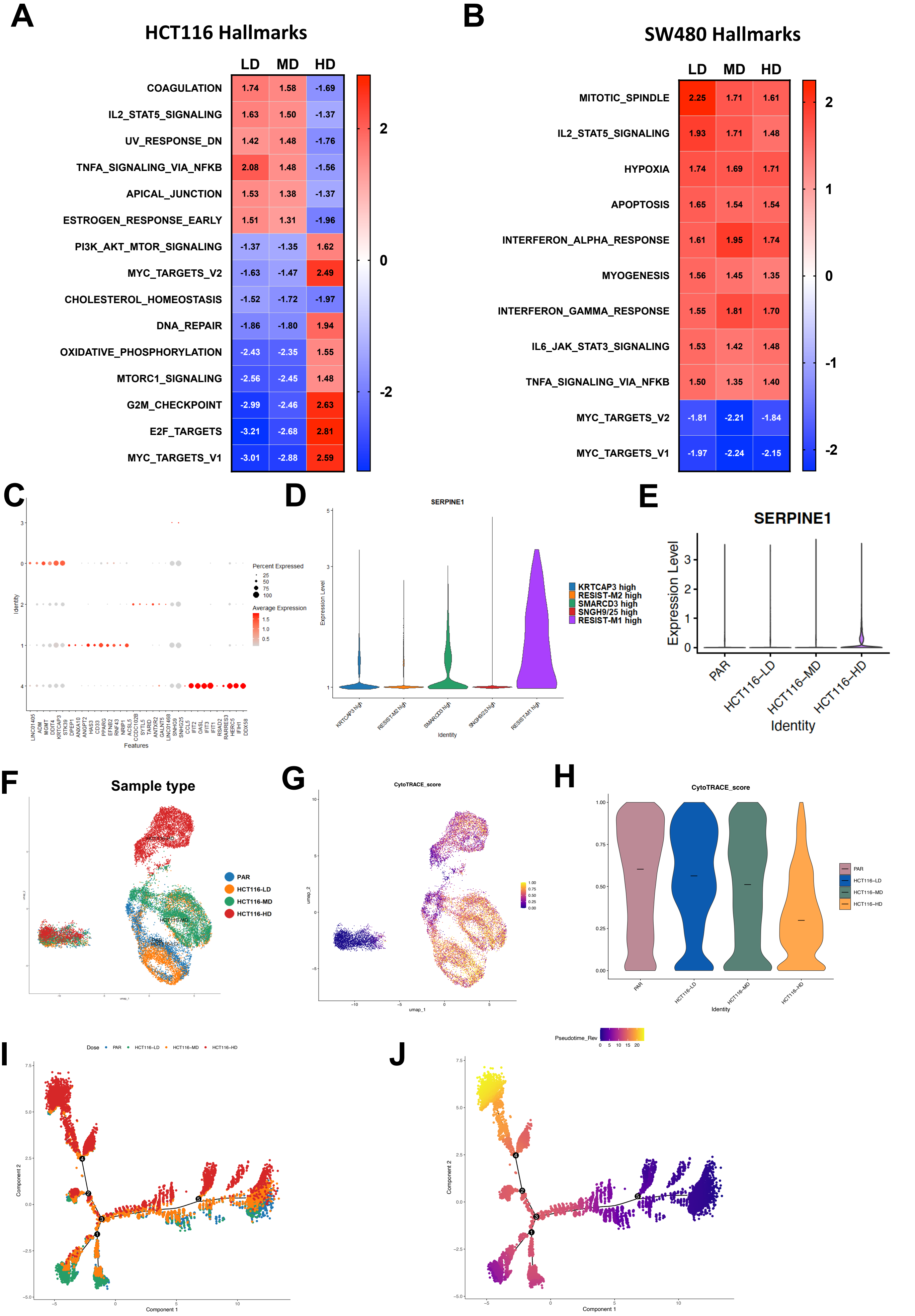

**Supplementary Figure 3. *SERPINE1* expression correlates with enhanced metastatic properties in human CRC patients and analyses of reported gene scores in our models**

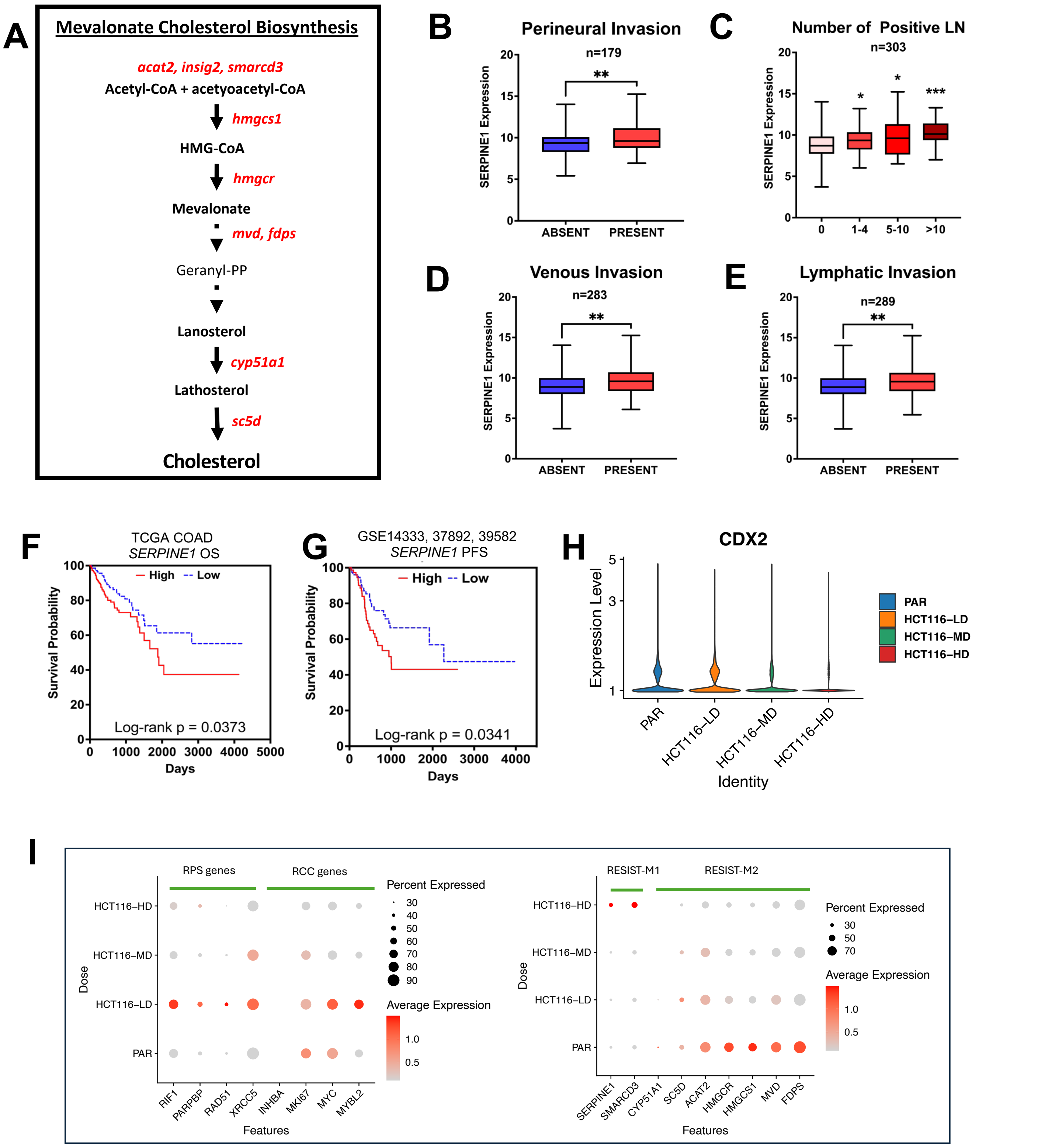

#### Supplementary Figure 4. RESIST-M expression correlates with poor prognosis in human CRC patients

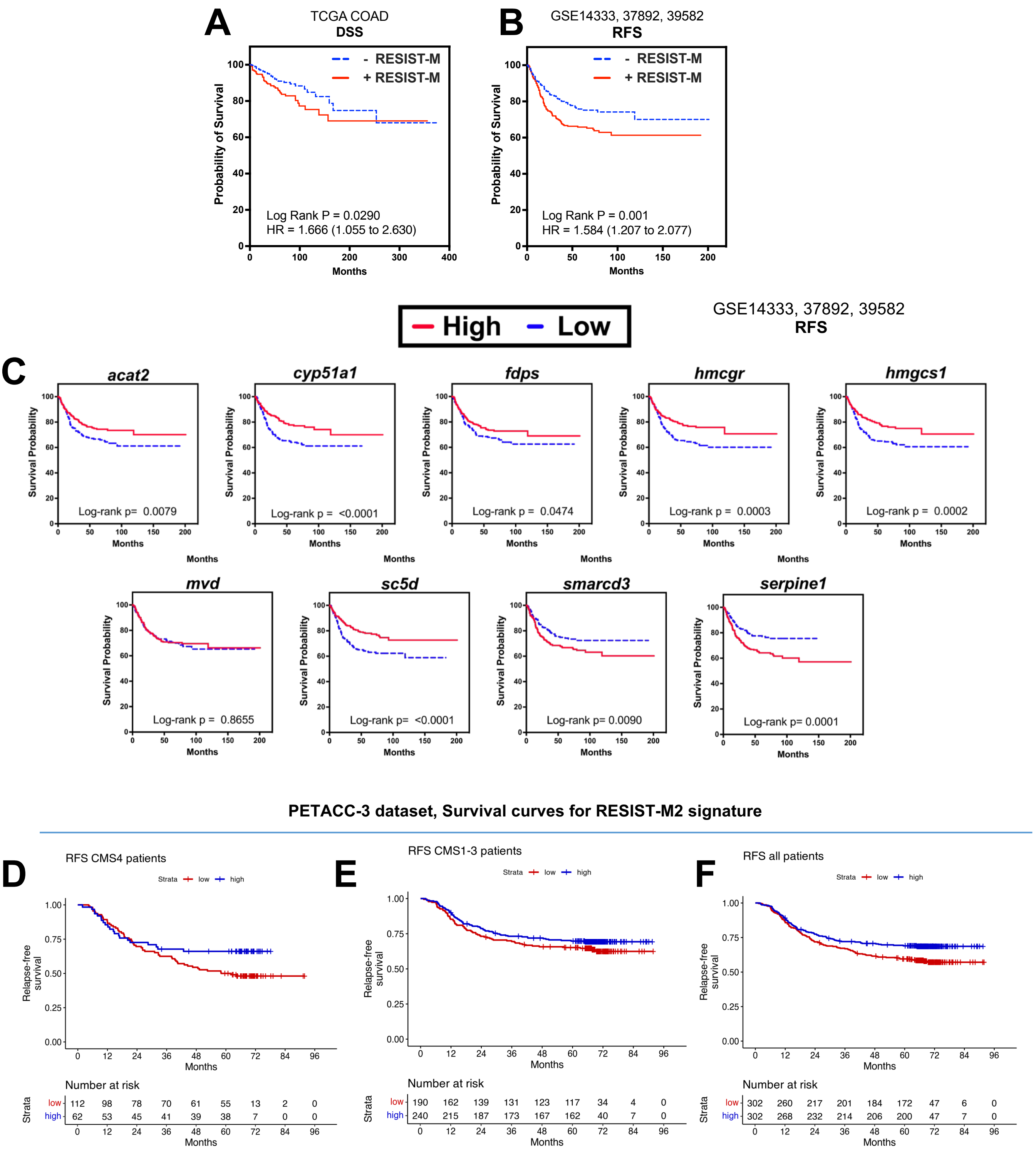

**Supplementary Figure 5. Therapeutic targeting of oxaliplatin resistant HCT116 and SW480 models**

**A**

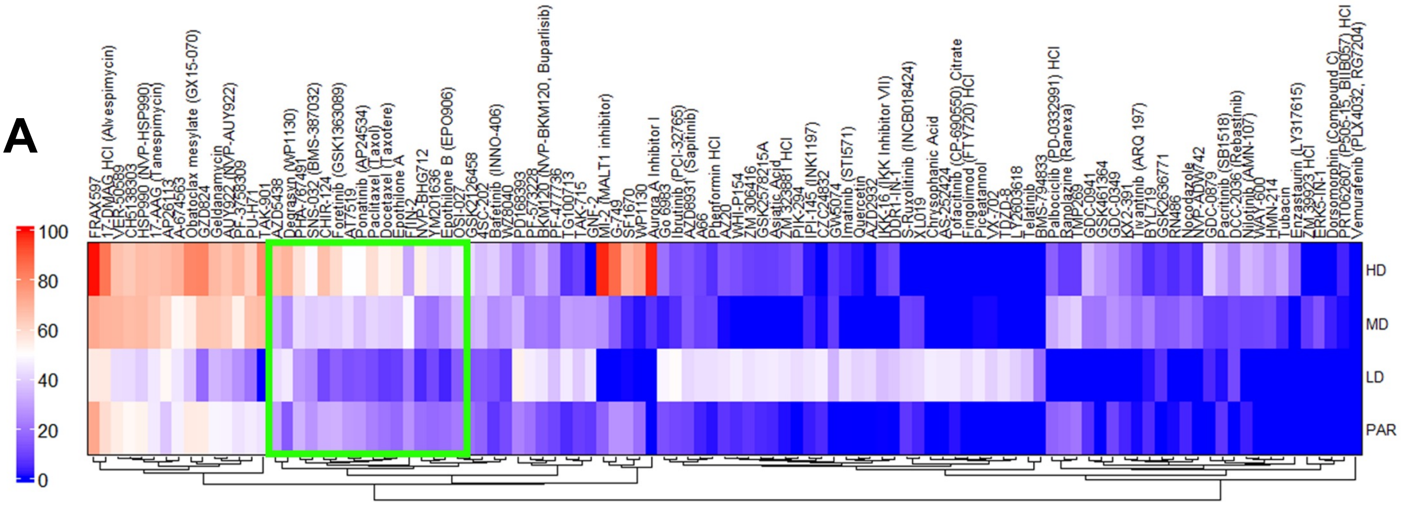


**B**

Inhibition score F.C. relative to PAR

| | LD | MD | HD | Target |
|---|---|---|---|---|
| AZD5438 | 1.41 | 1.10 | 2.19 | CDK1/2/9 |
| Degrasyn | 3.18 | 1.77 | 4.87 | USP5, UCH-L1, USP9x, USP14, UCH37, Bcr/Abl |
| PHA-767491 | 0.81 | 1.25 | 1.74 | Cdc7/CDK9 |
| SNS-032 | 0.89 | 1.44 | 1.81 | CDK2 |
| CHIR-124 | 0.32 | 1.25 | 2.04 | Chk1 |
| Foretinib | 0.48 | 1.19 | 1.80 | HGFR, VEGFR |
| AT7519 | 0.43 | 1.50 | 1.76 | CDK1, 2, 4, 6, 9 |
| Ponatinib | 0.47 | 1.31 | 1.81 | Abl, PDGFRα, VEGFR2, FGFR1, Src |
| Paclitaxel | 0.49 | 1.81 | 2.59 | Microtubule polymer stabilizer |
| Docetaxel | 0.38 | 1.97 | 2.64 | Inhibitor of depolymerisation of microtubules |
| Epothilone A | 0.48 | 2.17 | 3.07 | Paclitaxel-like microtubule-stabilizing agent |
| FIIN-2 | 0.73 | 1.83 | 1.17 | FGFR |
| NVP-BHG712 | 0.24 | 1.15 | 2.85 | EphB4 |
| YM201636 | 0.60 | 1.10 | 2.46 | PIKfyve |
| Epothilone B | 0.20 | 1.41 | 2.22 | Paclitaxel-like microtubule-stabilizing agent |
| OSI-027 | 0.68 | 1.58 | 2.12 | mTORC1/2 |

Color scale: 0 1 2 3 4

**Table 1. qPCR primers**

| Gene | Oligo Sequence (5' – 3') |
|---|---|
| SERPINE1_F | GCAACGTGGTTTTCTCACCC |
| SERPINE1_R | CTCTAGGGGCTTCCTGAGGT |
| VIMENTIN_F | TCTCTGAGGCTGCCAACCG |
| VIMENTIN_R | CGAAGGTGACGAGCCATTTCC |
| ACAT2_F | GCCTTGCAGTCCAGTCAATA |
| ACAT2_R | CTCAAGTAAGCCAAGTGAGGAG |
| HMGCS1_F | TATAGCTCTAGGTGTGCTCCTG |
| HMGCS1_R | CCTCATCCACACCTCCAATAAC |
| HMGCR_F | TTGCTTGCCGAGCCTAAT |
| HMGCR_R | GTTTCAGTCACCAACCTCCT |
| MVD_F | CCCAATGCCGTGATCTTCA |
| MVD_R | TCAGAAACGTGTCTCCATTCG |
| CYP51A1_F | CTCATCGCTCTTGCCAAATAAG |
| CYP51A1_R | TCCATCCTCACACACACAATAA |
| INSIG2_F | CTGATGCACAAGGGCAAGAT |
| INSIG2_R | GACACATCCTCTCACTCACTCC |
| SMARCD3_F | GATCAGTGCTCTGGACAGTAAG |
| SMARCD3_R | CTCTGGAGAAGCTTAGCATGAA |
| SC5DL_F | TCACTTTGTGGGATAGGATTGG |
| SC5DL_R | CGCTTTCCCTCTGTCATCTC |
| BACTIN_F | CACCATTGGCAATGAGCGGTTC |
| BACTIN_R | AGGTCTTTGCGGATGTCCACGT |
| GAPDH_F | AAGGGCATCCTGGGCTACACTGAG |
| GAPDH_R | GAAATGAGCTTGACAAAGTGGTCGTT |

**Table 2. Antibodies**

| Antibody | Manufacturer | Cat # | Dilution |
|---|---|---|---|
| PAI-1 | RnD Systems | MAB1786 | 1:500 |
| SMAD 2/3 | CST | 8685 | 1:1000 |
| p-SMAD2 (S465) | abcam | ab216482 | 1:1000 |
| p-SMAD3 (S434 + S425) | abcam | ab52903 | 1:1000 |
| SMAD4 | abcam | ab40759 | 1:1000 |
| Pan-ERK | CST | 4695 | 1:1000 |
| p-ERK 1/2 | Thermo | PA1-4607 | 1:1000 |
| Clathrin | abcam | ab21679 | 1:1000 |
| TGF-β RII | CST | 79424 | 1:1000 |
| GAPDH | Proteintech | 10494-1-AP | 1:5000 |
| Caveolin-1 | abcam | ab2910 | 1:1000 |
| MEK 1/2 | CST | 4694 | 1:1000 |
| p-MEK 1/2 | CST | 2338 | 1:1000 |
| p-AKT | CST | 4060 | 1:1000 |
| Histone H3 | CST | 4499 | 1:1000 |
| Cleaved- Caspase 3 | CST | 9661 | 1:1000 |
| Anti-Ms 680RD | LI-COR | 926-68070 | 1:10 000 |
| Anti-Rb 680RD | LI-COR | 926-68071 | 1:10 000 |
| Anti-Ms 800CW | LI-COR | 926-32210 | 1:10 000 |
| Anti-Rb 800 CW | LI-COR | 926-32211 | 1:10 000 |
| Anti-Rb HRP | CST | 7074 | 1:10 000 |
| Anti-Ms HRP | CST | 7076 | 1:10 000 |

#### Table 3. shRNA sequences

| Name | Oligo Sequence (5' – 3') | Notes |
|---|---|---|
| shControl_F | CCGGTCTCGCTTGGGCGAGAGTAAGCTCGAGCTTACTCTCGCCCAAGCGAGATTTTTG | |
| shControl_R | AATTCAAAAATCTCGCTTGGGCGAGAGTAAGCTCGAGCTTACTCTCGCCCAAGCGAGA | |
| PAI1 sh3 | CCGGCAGACAGTTTCAGGCTGACTTCTCGAGAAGTCAGCCTGAAACTGTCTGTTTTTG | TRCN0000052271, NM_000602.1-1041s1c1 |
| PAI1 sh5_F | CCGGGCTGACTTCACGAGTCTTTCACTCGAGTGAAAGACTCGTGAAGTCAGCTTTTTG | |
| PAI1 sh5_R | AATTCAAAAAGCTGACTTCACGAGTCTTTCACTCGAGTGAAAGACTCGTGAAGTCAGC | |
